# Reconstructing the emergence of the human chorion via HIPPO-mediated trophoblast induction

**DOI:** 10.1101/2025.11.18.688276

**Authors:** Miaoci Zhang, Ryan L. Lim, Alice H. Reis, Walter Piszker, Wallis W. Boyd, Athena Pagon, Aman Mahajan, Linlin Wu, Cheng Zhao, Sophie Petropoulos, Carlotta Ronda, Mijo Simunovic

**Author notes:** Correspondence to Mijo Simunovic.

## Abstract

The first lineage decision in the mammalian blastocyst commits outer cells to the trophectoderm and initiates the trajectory that gives rise to the placental chorion. The molecular sequence that unfolds downstream of HIPPO pathway inactivation, linking human trophectoderm specification to the early organization of the chorion, has remained unknown. Here, we establish a developmentally informed model that leverages HIPPO pathway modulation to induce the native trophectoderm trajectory in the absence of exogenous BMP or WNT signaling. We first transiently reset primed human pluripotent stem cells into a trophectoderm-competent ground state, followed by LATS kinase inhibition to set the trajectory in motion. To benchmark fidelity, we built an embryo-chorion single-cell reference integrating published early human and placental transcriptomes and applied a computational stage-matching tool to align our cultures to natural development. Stage matching revealed an ordered progression along the trophectoderm trajectory from early TE to post-implantation trophoblast. With extended culture, all major cell types of the nascent chorion emerged, encompassing both trophoblast and chorionic mesoderm lineages. Within the trophoblast, we identified proliferative and non-cycling villous cytotrophoblast, a columnar population connecting villous and extravillous domains, as well as syncytia and extravillous subtypes. When cultured in suspension, these lineages self-organized into three-dimensional organoids that recapitulated the stromal-epithelial architecture and proliferative-syncytial polarity of the emergent chorion. We identified CLDN6 as a defining surface marker of columnar trophoblast, the population that bridges villous and extravillous compartments. Prospective isolation of living CLDN6+ trophoblast revealed their capacity to reacquire a proliferative villous state and, under directed cues, generate both syncytial and extravillous fates, confirming their proposed dual developmental potential within the chorion. Together, these findings establish a developmentally informed framework that connects human trophectoderm specification to the emergent chorion and provides a dynamic platform for investigating the earliest steps of placental specification and the origins of implantation disorders.

## Introduction

The first lineage decision in mammalian development occurs at the blastocyst stage, when outer blastomeres commit to the trophectoderm (TE). This epithelium mediates implantation and later gives rise to the trophoblast (TB) lineages, which form and regenerate the chorionic villi of the placenta^1^. The chorion marks the first organized placental structure, emerging by the end of the second week of human development when TBs form villous and extravillous compartments surrounding a mesodermal core. This stage establishes the foundation for villous architecture and the progenitor populations that sustain placental growth throughout gestation. Defects in early chorion formation contribute to implantation failure and placental insufficiency disorders such as preeclampsia and intrauterine growth restriction, which together account for substantial maternal and neonatal morbidity worldwide^2,3^. Yet despite their clinical importance, the cellular and molecular events linking TE initiation and guide its progression toward nascent chorion remain poorly defined.

Multiple strategies now enable robust derivation of human TB from pluripotent stem cells, such as protocols using BMP4 with varying inhibitors, or WNT-based media that can chemically convert naïve cells to TB stem cells^4–19^. A recent computational tool that stage matches single cell transcriptomes of *in vitro* data against an integrated embryonic reference indicated that existing approaches recapitulate TB lineages with varying fidelity^20^. Most current methods rely on exogenous morphogen combinations, often following purification steps via fluorescence-based sorting or multiple passaging to isolate and expand cells with proliferative cytotrophoblast (CTB) identity in TB stem cell media. What remains missing is a developmentally informed, experimentally tractable framework that traces the native transition from an early embryonic, TE-competent state to the establishment of the nascent chorion, the basic building block of the placenta.

Work in the mouse established that Hippo signaling pathway is the primary mammalian regulator of the first cell fate decision. In outer cells, polarization inhibits LATS1/2 kinase activity, allowing nuclear accumulation of the transcriptional coactivator YAP and the activation of TEAD-dependent transcription^21–24^. Genetic loss of *Tead4* prevents TE formation and causes preimplantation arrest, while genetic disruption of *Lats1/2* or other pathway components (*Nf2, Amot*) leads to ectopic induction of TE genes^22– 26^. Despite species-specific differences in early mammalian development, comparative embryo studies confirm the evolutionary conservation of the Hippo-dependent mechanism across several phylogenetically distinct mammalian blastocysts^27,28^. Understanding how HIPPO suppression drives the sequential emergence of TB states toward the nascent chorion provides a mechanistic foundation for reconstructing early human placental development.

To model the emergence of TE in a way that reflects embryonic development, the most appropriate approach is to mimic this LATS-dependent HIPPO pathway inactivation. Experiments across a wide range of mammalian embryos, namely cows, humans, guinea pigs and other rodents, has shown that culturing embryos with a LATS kinase inhibitor ectopically activates TE genes in the ICM^27–29^. Work on human embryo models has shown that lysophosphatidic acid (an upstream inhibitor of LATS kinases) contributes to the proper establishment of TE identity in blastoids^30^. YAP over-expression can also yield cells with TE identity^10,31^. Collectively, these studies implicate LATS kinase inhibition as a likely, developmentally faithful trigger for activation of the TE gene program. Building on this principle, we set out to establish a framework that leverages targeted manipulation of the HIPPO-LATS-YAP axis to capture the progression of human TB development from a TE-competent ground state to the emergence of the nascent chorion.

## Results

### Establishing a transient window of TE competence in primed hESCs

Prior to lineage bifurcation, cells in the embryo possess the competence to respond to HIPPO signaling and initiate the TE trajectory, ultimately leading to all the TB lineages of the placental chorion. Naïve hESCs most closely resemble this state, but maintaining them long-term poses technical and biological challenges, including culture instability and imprinting loss, depending on the precise culture conditions^32–34^. As a more practical approach, we established a strategy designed to transiently mimic the transcriptional and epigenetic state of naïve cells (Fig. 1a). To approximate the chromatin and DNA methylation characteristics of the early embryo, we supplemented naïve media with two small molecules: GSK34834862, a selective DNMT1 degrader (DNMT1i), and DZNep, an indirect EZH2 inhibitor (EZH2i). GSK34834862 has previously been shown to efficiently and reversibly induce global demethylation of murine ESCs^35^, and we confirmed it eliminates DNMT1 protein in hESCs within 24h (Fig. S1a). DZNep reduces H3K27me3 in hESCs^35–37^. Furthermore, in naïve hESCs, EZH2/PRC2 inhibition selectively derepresses bivalent promoters at genes that prime naïve cells for TE competence^37^. After testing several naïve media, we settled on a modified human enhanced stem cell medium (HENSM)^9^ supplemented with the two epigenetic inhibitors, which in our hands yielded the most robust transient naïve reset with high cell viability.

**Figure 1:**
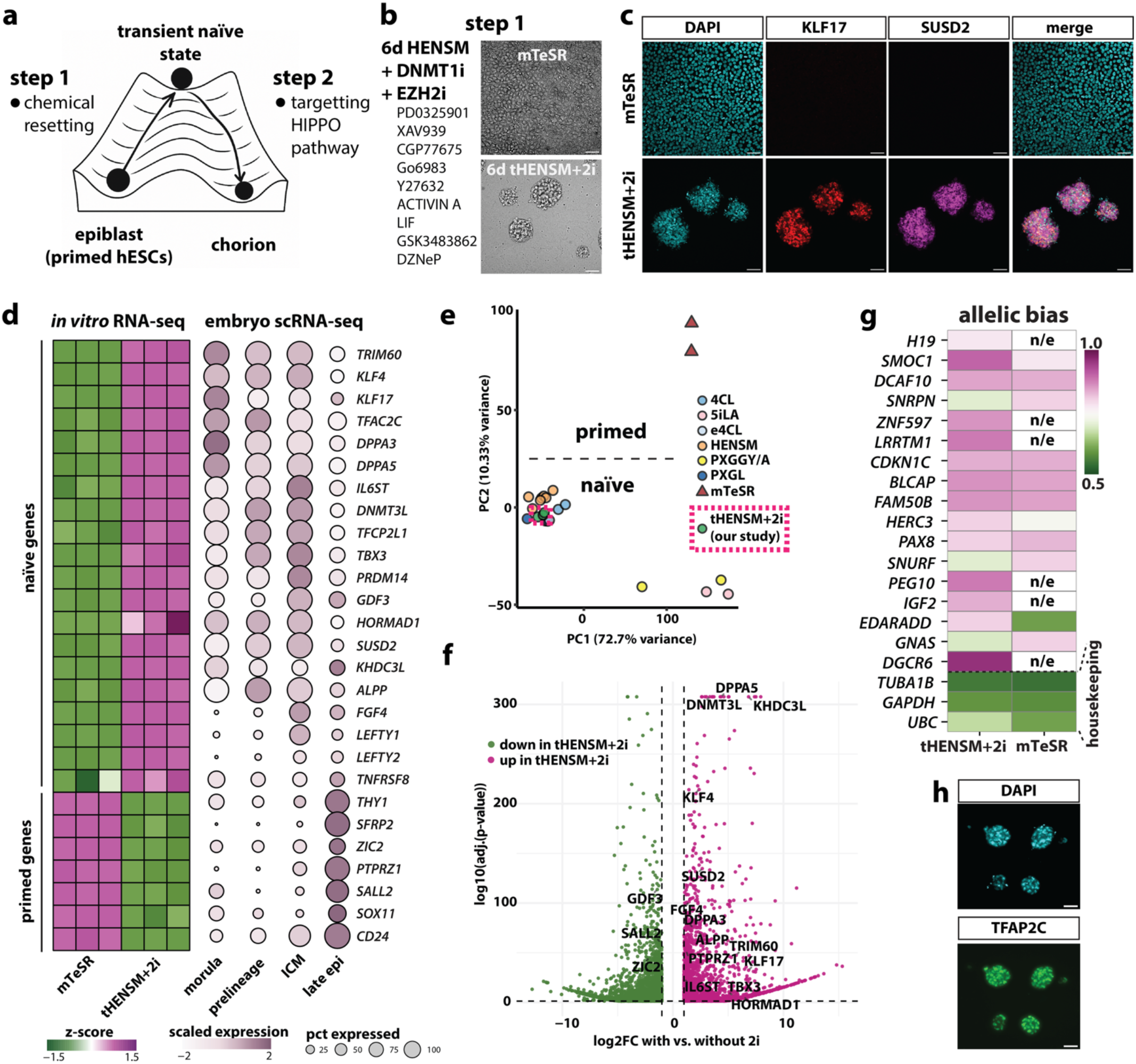
Transient resetting to a prelineage state. (a) Schematic of the two-step strategy: (1) transient naïve resetting of primed hESCs; (2) subsequent induction of the TE program via HIPPO modulation. (b) Phase contrast images of hESCs cultured in HENSM + DNMT1i and EZH2i showing dome-like naïve morphology within six days. (c) IF for naïve hESC markers. (d) Bulk RNA-seq comparison between mTeSR and tHENSM+2i conditions and the single-cell human embryo reference^20^; n = 3 for each condition. (e) PCA positioning tHENSM+2i relative to published naïve and primed data. (f) Volcano plot of differential expression between bulk-seq HENSM+2i and HENSM highlighting upregulation of naïve hESC genes and a downregulation of primed genes in the epigenetic drug treated condition. (g) Allele-specific expression from bulk RNA-seq showing preserved monoallelic expression across imprinted loci, compared to primed control. n/e, not expressed (h) IF for TFAP2C in day 6 HENSM+2i aggregates. All scale bars, 50 µm.

Once transitioned into HENSM+2i, hESCs progressively upregulated naïve-associated genes (Fig. S1b). By day six, they compacted into characteristic dome-shaped morphologies (Fig. 1b) uniformly expressing naïve markers SUSD2 and KLF17^38^, while still expressing broad pluripotency markers (Fig. 1c, S1c). We refer to this transient condition as tHENSM+2i. We tested three hESC lines (two XX and one XY), all showing comparable morphology and naïve gene expression in tHENSM+2i condition (Fig S1e,f).

We performed bulk RNA sequencing of the tHENSM+2i and mTeSR conditions and compared their transcriptomes with single-cell RNA transcriptomes of human embryos, pseudobulked by stage. tHENSM+2i showed a robust downregulation of late-epiblast-associated genes (*THY1, SFRP2, ZIC2, DUSP6*) and induction of genes whose expression is shared among the morula, prelineage, and ICM, consistent with naïve state (*KLF2, KLF4, KLF17, IL6ST, DPPA3, DPPA5*) (Fig. 1d). Overall, comparison of the transcriptomes with the 50 most differentially expressed genes from distinct embryo stages further confirmed that the mTeSR condition more closely resembled the late epiblast, whereas tHENSM+2i more closely resembled earlier embryonic states (Fig. S1g). Principal component analysis confirmed that tHENSM+2i clustered closely with published long-term naïve cultures along the first two principal components and distinctly away from primed hESCs, confirming that brief resetting is sufficient to invoke a global naïve transcriptional state (Fig. 1e).

We also assessed the influence of epigenetic inhibitors on the bulk transcriptome in tHENSM+2i condition, and found that they further upregulated naïve-associated genes, including *DPPA5, DNMT3L, KLF4, DPPA3* and others (Fig. 1f). Because prolonged naïve culture is known to erase imprinted genes^33^, we calculated allele-specific RNA expression from bulk RNA-seq and observed robust preservation of monoallelic expression across many known placental imprinted genes, when compared to primed hESCs (Fig. 1g). Transient resetting also induced expression of TFAP2C protein (Fig. 1h). In the human embryo, TFAP2C is expressed prior to polarization and its expression is critical in establishing a bipotent transcriptional state through co-activation of both TE and ICM regulators^39^. Consistent with global epigenetic resetting, LINE-1 elements, markers broadly correlated with DNA hypomethylation, were prominently upregulated in tHENSM+2i (Fig. S1h).

Together, these findings show that transient resetting with tHENSM+2i rapidly induces morphological, transcriptional, and epigenetic features of the naïve, TE-competent state, while preserving imprinting integrity.

### HIPPO pathway modulation induces a high-fidelity TE program independently of BMP and WNT

TE specification in the embryo is initiated by suppression of the HIPPO pathway, which promotes nuclear localization of YAP and activation of a TEAD-dependent transcriptional program, as detailed in the introduction (Fig. 2a). To model this transition *in vitro*, we used a selective LATS inhibitor TDI-011536 (TDI), a more potent derivative of a small molecule TRULI, which was previously shown to in-duce nuclear YAP shuttling in hESCs^40^. Specifically, after a transient reset in tHENSM+2i, we applied TDI±A83-01 for the first six days, followed by TDI alone for additional six days, in otherwise neutral media (Fig. 2b). Immunofluorescence (IF) revealed strong nuclear accumulation of YAP and a reduction of phosphorylated YAP (pYAP), consistent with HIPPO inhibition (Fig. 2c). To monitor TE induction, we engineered a CRISPR/Cas9 T2A-tdTomato knock-in reporter in the *GATA3* locus (Fig. S2a). It allowed us to record a progressive GATA3 activation across the culture period, reaching near-uniform expression by day six. It took place with or without the A83-01 pulse (Figs. 2d and S2a), although there was a slightly more accelerated GATA3 response under A83-01 pulse (Fig. S2a).

**Figure 2:**
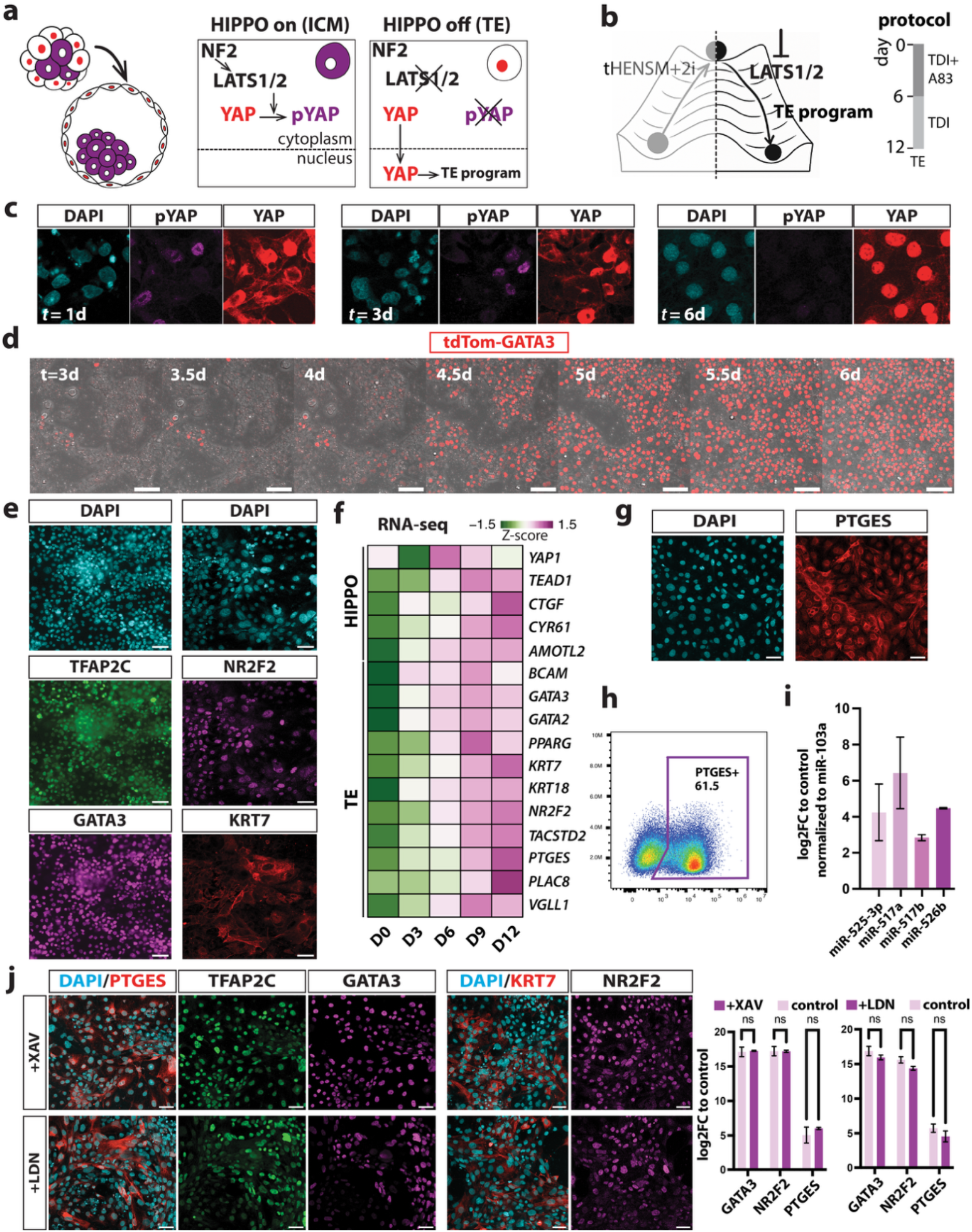
Induction of the TE program via HIPPO modulation independently of BMP or WNT. (a) Schematic summarizing the role of HIPPO-LATS1/2-YAP axis in inducing the TE program in a eutherian embryo. (b) Experimental design for HIPPO-dependent TE induction following the transient prelineage resetting. (c) IF of pYAP and total YAP showing progressive nuclear YAP accumulation following LATS inhibition. (d) Time-lapse imaging using the tdTom-GATA3 reporter showing high-yield and timely induction. (e) IF of TE-associated markers on day 12 of differentiation. (f) Bulk RNA-seq at regular time intervals showing the upregulation of HIPPO and TE associated genes (n=3, except n=2 for day six). (g,h) IF and flow cytometry showing broad expression of PTGES (n = 2 pooled). (i) qPCR of primate-specific C19MC microRNA cluster on day 12 compared to primed hESCs (n = 3) (j) IF and qPCR analysis of XAV939 and LDN193189-treated cultures, showing no change in TE marker expression (n = 3, two-way ANOVA, P < 0.05, ns = not significant). Error bars represent mean ± s.d. Scale bars, 50µm (c,e,g,j) or 100µm (d).

Multiple TE-associated proteins, including TFAP2C, NR2F2, GATA3, and KRT7, progressively accumulated over the 12-day induction (Figs. 2e and S2b). On day 12, there were no detectable epiblast-like cells (Fig. S2c). Bulk RNA-seq profiling on days 0, 3, 6, 9, and 12 showed induction of HIPPO/YAP targets (*YAP1, TEAD1, CTGF, CYR61, AMOTL2*) and a strong enrichment of *YAP*/*TAZ* target genes by day 12 (Figs. 2f and S2d). By this point, transcriptomic profiling showed the expression of all known canonical TE genes (e.g., *GATA3, GATA2, PPARG, KRT7, KRT18, NR2F2, TACSTD2, PTGES, PLAC8, VGLL1*) ^19,41^(Fig. 2f). To further benchmark lineage fidelity, we applied the 50 most differentially expressed genes in the TE cluster from the integrated single cell embryonic reference^20^ to our bulk RNA-seq time course. TE-cluster genes became progressively enriched from days three to 12 (Fig. S2e). By contrast, our model showed much lower correspondence to amnion-cluster genes compared to the published transcriptome of BMP4/A83-01/PD173074-treated primed cells^19^ (Fig. S2e).

We observed additional hallmarks of authentic TB. Implantation-associated enzyme prostaglandin E synthase (PTGES) was detected in >60% of cells by flow cytometry and confirmed to be broadly expressed at the protein level by IF (Fig. 2g,h). The primate-specific C19MC microRNA cluster, which is essential for TB development and is silenced in primed hESCs^42^, was indeed strongly induced (Fig. 2i). Bisulfite sequencing showed hypomethylation of the *ELF5* promoter compared to primed cells, another epigenetic defining features of TB identity^41^ (Fig. S2f).

Finally, we were curious if HIPPO-driven induction requires input from WNT or BMP, given that most *in vitro* TB induction or TB stem cell protocols depend on exogenous BMP or WNT stimulation. Interestingly, inhibition of WNT with XAV939 or BMP with LDN193189 had no significant difference in the gene or protein expression of canonical TE targets, including GATA3, NR2F2, and PTGES (Fig. 2j).

Taken together, HIPPO suppression alone is sufficient to drive a robust TE program in transiently reset hESCs, independently of BMP or WNT input.

### Transcriptional stage matching reveals progressive differentiation along the TE trajectory

We next sought to define the temporal sequence of transcriptional and lineage transitions that unfolds downstream of HIPPO suppression and align it with the embryonic trajectory. First, we extended culture beyond day 12 by removing TDI and monitoring progression toward the CTB lineage with ITGA6 protein expression^4^. We found that cells spontaneously express ITGA6 in basal medium (see Methods), and expression increased modestly when cultures were supplemented with CHIR99021 and EGF, factors previously shown to be supportive of TE maturation and TB stem cell maintenance^4,19^ (Fig. S3a).

We collected cells on days 6, 12, and 18 of differentiation, retaining ∼3000 cells per time point after quality control (see Methods). To stage match the HIPPO-mediated TB trajectory with embryonic development, we projected our single cell transcriptomes onto the UMAP space of an early human embryonic reference, following a recently published method^20^. The integrated reference resolves the entire TB trajectory from the first cell fate choice in the human embryo to embryonic day 14 (Fig. 3a).

**Figure 3:**
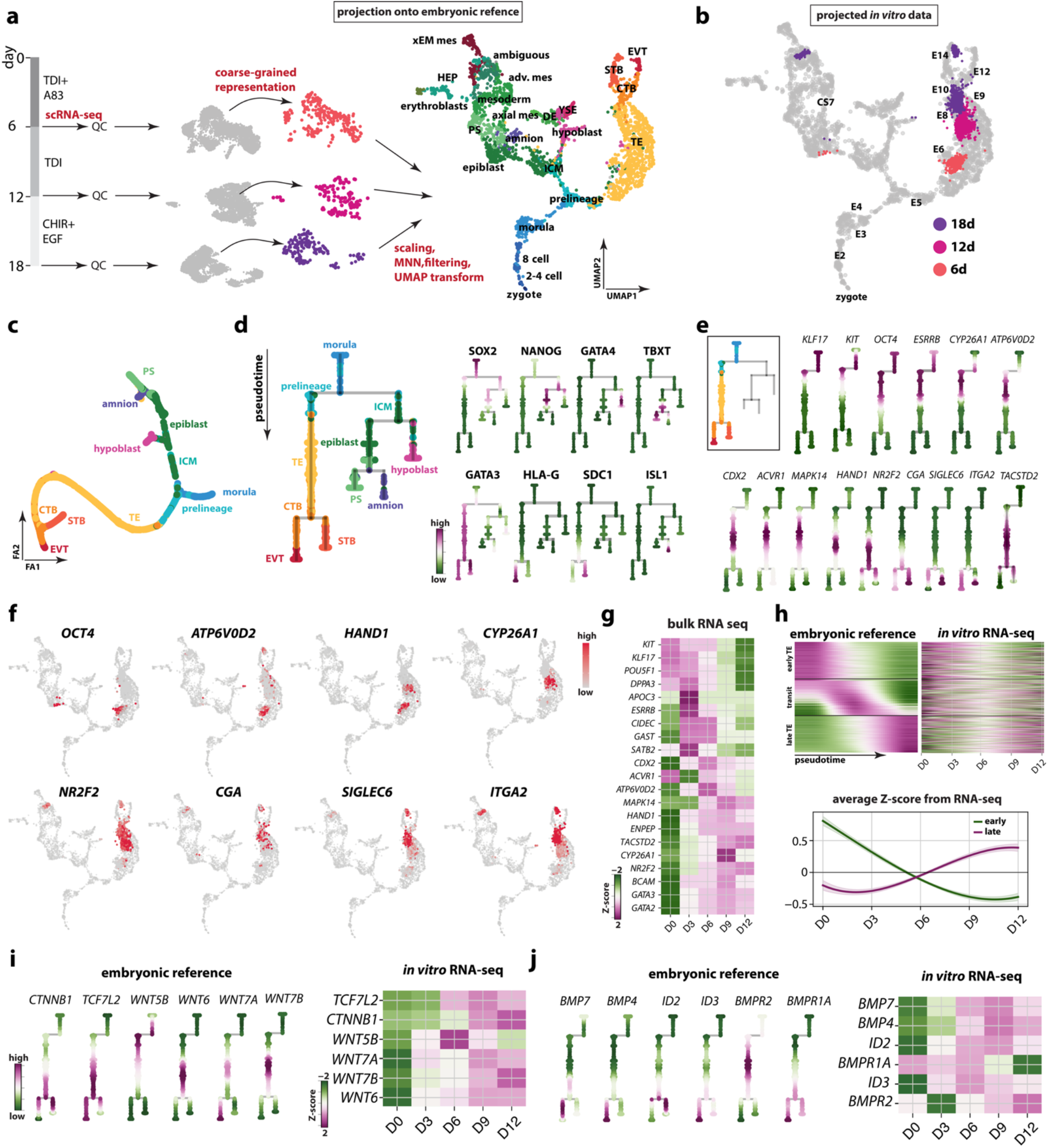
Temporal gene expression dynamics downstream of LATS inhibition follow the embryonic TB trajectory. (a) Projection of single-cell transcriptomes collected on days 6, 12, and 18 of differentiation onto the integrated embryonic reference (zygote to CS7) to stage match in vitro samples to developmental time in vivo. The reference UMAP is annotated as in Ref. 20. (b) Projection of in vitro datasets shows sequential progression along the TB lineage. (c) ForceAtlas2 (FA2) layout of the embryonic reference generated in diffusion space with Palantir. Embryonic reference is subset as described in the text. (d) Pseudotime trajectory derived from the FA2 embedding using scFates and represented as a dendrogram, displaying the expression of lineage-representative markers. (e) Expression dynamics of genes illustrating successive phases of TB differentiation. (f) Expression of selected TB genes mapped onto the integrated embryonic reference. The embryonic reference is shown in gray, while color intensity reflects gene expression in the projected in vitro cells. (g) Bulk RNA-seq at regular time intervals showing temporal dynamics of TE-associated genes (n = 3, except day 6 = 2). (h) Left: heatmap showing genes ordered along the TB branch of the pseudotime trajectory in the embryonic reference and grouped into early, intermediate, and late modules. Displayed only 10% of each module for clarity. Right: the same gene order was applied to the in vitro bulk RNA-seq time course. Below: averaged Z-scores from bulk RNA-seq illustrate the reciprocal dynamics of early and late modules during differentiation. (i-j) WNT and BMP pathway activity show a similar expression dynamics in vitro to the embryonic reference.

In this projection approach, query single cell transcriptomes are coarse-grained into neighborhoods, then normalized, scaled, and matched to reference cells by mutual nearest neighbors, and finally placed into the stabilized reference UMAP (Fig. 3a)^20^. Projection of our *in vitro* transcriptomes resolved an ordered progression along the TB branch: day 6 aligned near pre-implantation TE, day 12 at peri-implantation TE, and day 18 distributed along the axis connecting CTB, syncytiotrophoblast (STB), and extravillous TB (EVT) (Fig. 3b).

To delineate the cell fate choices along the trajectory, we clustered the combined projected *in vitro* datasets preserving the UMAP embedding of the reference, which revealed three distinct regions (Fig. S3b): (i) *OCT4*+ early/transitional cluster, consistent with the residual *OCT4* expression at prelineage to TE transition in the embryo (Fig S3c), (ii) *KRT7*+ *GATA3*+ pan-TB clusters, which we subdivided into pre-, peri-, post-implantation, and CTB derivatives, based on how closely they projected onto the equivalent stages on the embryonic reference (Fig. S3d), and (iii) *POSTN*+ *COL3A1*+ stromal cluster that projected adjacent to the extraembryonic mesoderm in the reference (Fig. S3e). Of note, we resolve the precise lineage diversity and assess their similarity to the nascent chorion in the next section; here, we focus on characterizing the dynamics of the *in vitro* trajectory.

To quantify serial order along the TB trajectory, we subset the embryonic reference to exclude all stages prior to morula and, from the CS7 dataset (which did not include the chorion), retained only the primitive streak and amnion clusters. We constructed a force-directed embedding in FA space (Fig. 3c) then applied *scFates* to the FA embedding^43^, and fitted gene expression profiles along it. We represent the trajectory with a dendrogram throughout this section to provide a framework for continuous transcriptional dynamics (Fig. 3d).

Ordered gene expression demarcated transitions along the TB lineage (Fig. 3e). The transition from prelineage into TE was marked by *KIT* and *KLF17*, together with tapering of *OCT4* and *ESRRB*. These genes were also transiently expressed in the first stages of differentiation *in vitro* (Fig. 3f,g). The expression of *CDX2, HAND1, ATP6V0D2*, and *CYP26A1* is seen in early and mural TE in the human embryo, known to play roles in blastocoel formation and retinoic metabolism^44^. The same genes showed earlier expression in our model as well. *NR2F2*, a gene marking polar TE across mammals^29,45^, appeared subsequently in the pseudotime trajectory, showing a similar expression dynamic in our differentiation (Fig. 3e-g). Later in the trajectory, genes including *TACSTD2, CGA, SIGLEC6*, and *ITGA2*, all associated with TE maturation into CTB, and chorionic TB induction, became progressively enriched, mirroring the temporal logic observed *in vivo*.

Relevant long non-coding transcripts and small-RNA regulators also followed characteristic lineage-asymmetric patterns along the trajectory, reflecting imprinting and small-RNA control of early lineage segregation. As in the human embryo, *XIST* remained active, consistent with preserved X-chromosome dosage compensation. The maternally expressed transcripts *MEG3* and *MEG8* from the *DLK1-DIO3* locus, markers of intact maternal imprinting, and hosts for the C14MC miRNA cluster, were abundant at early stages and declined with differentiation, consistent with the physiological attenuation of maternal non-coding RNA expression as the TE trajectory unfolds (Fig. S3f)^42,46^. *DICER1* remained broadly expressed throughout the trajectory, indicating active machinery required for small-RNA biogenesis^42,46^ (Fig. S3f).

Looking broader, we grouped all genes in the embryo reference into early, intermediate, and late bins based on their fitted expression along the pseudotime trajectory, displaying a subset of each group for clarity (Fig 3h, left; Fig. S3g). Applying the same gene order to our bulk RNA-seq data exposed a coordinated global switch from early to late transcriptional modules, with early modules declining and late ones rising starting around day six of differentiation (Fig. 3h, right and bottom; Fig. S3g).

Finally, we asked whether endogenous WNT or BMP signaling appears downstream of HIPPO. We first characterized the expression of downstream response genes of these pathways in the embryo, finding *CTNNB1, TCF* factors, and *ID* genes undetected at the prelineage to TE transition, indicating that WNT and BMP signaling are likely not active during initial TE induction (Fig. 3i,j). Interestingly however, various ligands appeared downstream with distinct timing: a noncanonical *WNT5B* was the prominent WNT ligand expressed first in the TB trajectory, whereas canonical *WNT6, WNT7A, WNT7B* were expressed later. BMP ligands associated with placenta biology such as *BMP4* and *BMP7* were also expressed later in the TE trajectory, along with their downstream targets *ID2* and *ID3*. Our *in vitro* cultures mirrored this pattern, showing early *WNT5B* expression and later induction of the other WNT and BMP ligands (Fig. 3i,j). These observations corroborate our finding that HIPPO suppression alone initiates TE specification, although they show an interesting pattern of autocrine WNT and BMP activity emerging downstream accompanying chorionic emergence.

### Induction and self-organization into emergent chorion

Once CTB is established in the embryo, it contributes to the primary villus, in which villous CTBs (VCTs) maintain the epithelial surface, fuse to form STB, and, in anchoring villi, extend into cell columns whose proximal cells act as progenitors for the EVT lineages. The secondary villus forms as extraembryonic mesoderm fills the villous core forming chorionic stroma (Fig. 4a). Having established that day-18 cultures align with the most advanced TB states on the embryonic reference, we wondered to what extend they recapitulate the cellular composition of the nascent chorion.

**Figure 4:**
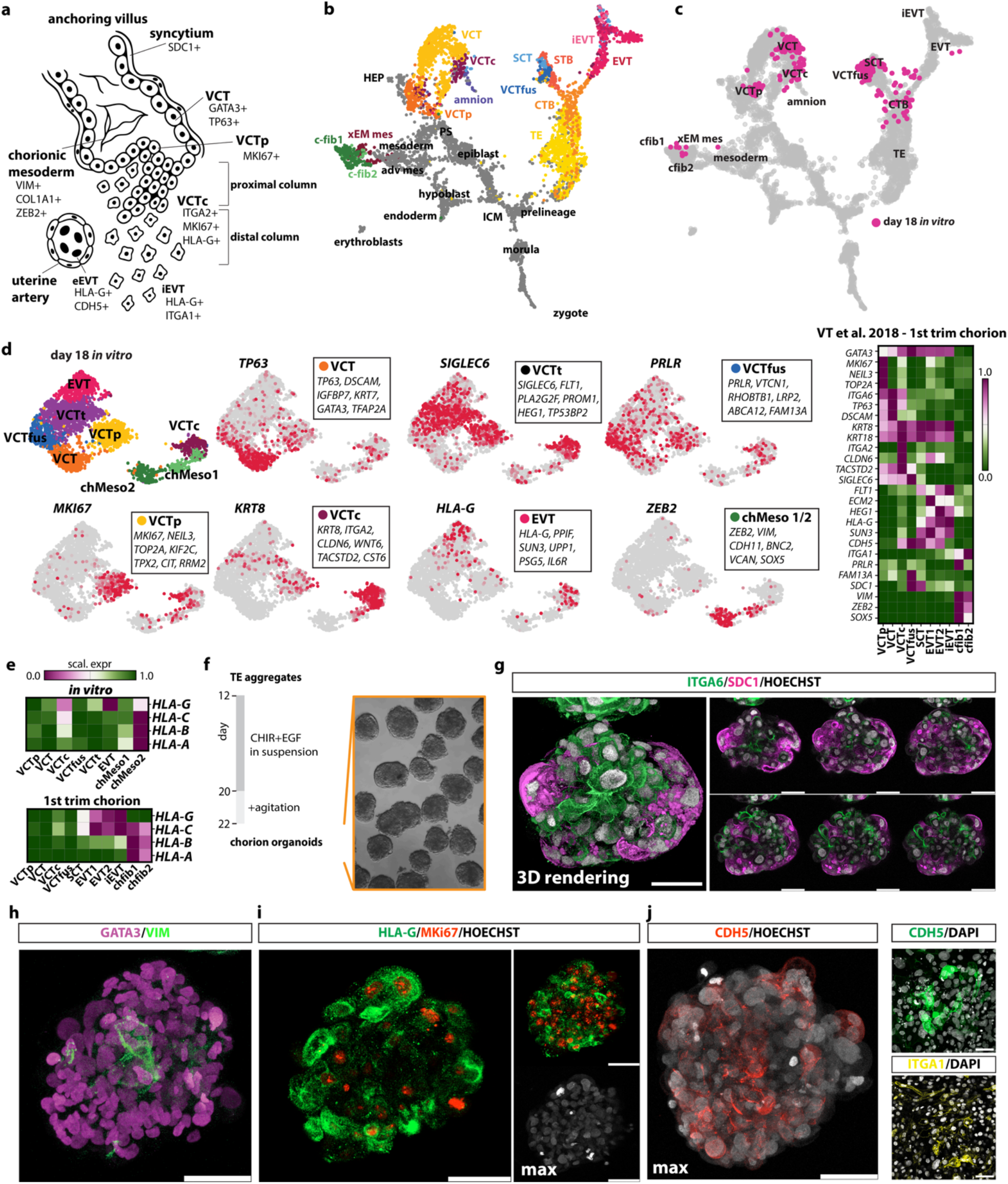
The emergence of chorionic lineages and chorionic organoids in the differentiation trajectory. (a) Schematic of the anchoring chorionic villus showing spatial arrangement and marker profiles of known chorionic subtypes. VCT=villous cytotrophoblast; VCTp=proliferative VCT; VCTc=column VCT; iEVT=interstitial EVT; eEVT=endovascular EVT. (b) UMAP of the extended integrated embryonic-placental reference map spanning zygote to first-trimester villi. Chorionic lineages are color-coded, following original annotations. (c) Projection of day-18 in vitro single-cell transcriptomes onto the extended reference showing representation across all major chorionic lineages. (d) UMAP of day-18 single-cell profile and cluster annotations. Right: scaled expression of lineage-specific markers from the first-trimester chorion reference^47^. (e) Scaled expression of HLA family genes compared with the first-trimester chorion reference. (f) Workflow of chorion organoid generation. (g-j) IF images of chorion organoids showing: correct polarity with SDC1+ syncytia overlying ITGA6+ VCT cells (g), stroma-TB organization (h), and the emergence of EVTs (i,j – subset in j right shows adherent culture on day 18). Further examples shown in Fig S6). Scale bars, 50 µm.

To resolve the chorionic subtypes in early development, we extended the embryonic reference with the published single-cell transcriptome of the first trimester placenta^47^, similarly to the reference extension described in the original projection tool^20^. Here, we included both TB and stromal lineages of the chorion, coarse-grained cells to avoid overrepresentation, then scaled, normalized and integrated with six embryonic transcriptomes spanning from zygote to CS7^16,45,48–51^. We preserved the annotations from the first trimester placenta study and carried over the annotations of the integrated embryonic reference^20^ (Fig 4b). The resulting extended map encompassed datasets from the zygote through first-trimester chorion, providing a unified framework for embryo-placenta comparison (Fig. 4b, Fig. S4a,b).

To test fidelity of the extended reference map, we projected two independent datasets not used in the integration: an embryonic day 10 (E10) human embryo we published previously^52^ and a published single nuclear sequencing dataset of first trimester placenta from the A. Moffett collection^53^. Both datasets projected onto the map matching their independently annotated stage and cell types (Fig. S4c). We then projected our day-18 *in vitro* datasets. Projection onto the extended embryo-placenta reference showed representation of all TB and stromal lineages of the chorionic villus, with a smaller portion of coarsegrained neighborhoods aligning with advanced EVTs (Fig. 4c). We also projected our day-12 alongside the day-18 dataset (Fig. S4d) which aligned similarly to the original embryonic reference with the majority of cells aligning near the peri-implantation TE, as further validation.

We next examined the cellular composition of our day-18 dataset in detail. Unsupervised clustering of single-cell profiles on a UMAP embedding resolved nine distinct groups of cells (Fig. 4d). We identified representative genes using differential expression analysis and compared them with transcriptional signatures from the first-trimester placenta reference (Fig. 4d). We distinguished two main VCT populations from the chorion: a proliferative *MKI67*+, *NEIL3*+, *TOP2A*+ subset (VCTp) and a *TP63*+, *ITGA6*+, *DSCAM*+ subset corresponding to non-cycling VCTs (Figs. 4d and S5). A separate cluster expressing *ITGA2, KRT8, CLDN6, WNT6*, and *TACSTD2* corresponded to the VCTs in the columns of anchoring villi. This cluster showed a gradient of *MKI67+* cells, consistent with proximal proliferative column TBs, versus *HLA-G*+ cells, consistent with differentiating distal column TBs progressing toward EVT. We therefore annotated it as columnar VCT (VCTc). A *GATA3*+, *ITGA6*−, *PRLR*+, *VTCN1*+ cluster matched fusion-competent VCTs (VCTfus), while an *HLA-G*+, *PPIF*+, *SUN3*+, *IL6R*+ cluster corresponded best to EVTs. In addition, we identified a *GATA3*+, *ITGA6*+, *SIGLEC6*+ cluster carrying a mixed transcriptional signature seen in both villous and extravillous subtypes in the placenta, such as *PLA2G2F, FLT1, HEG1*, and *TP53BP2*, and we annotated it as transitional VCT (VCTt). Finally, two clusters expressed mesenchymal signatures, including *VIM, ZEB2, SOX5*, and *VCAN*, consistent with the chorionic stroma and we annotated is as chorionic mesoderm (chMeso).

We also examined the distribution of class I HLA transcripts in the single cell transcriptomes. Namely, HLA-A/B/C were only robustly detected in chMeso cluster, with HLA-G in EVT and VCTc clusters, in line with their expression patterns in the first trimester chorion (Fig. 4e).

To visualize spatial organization and protein expression of the emerging chorionic lineages, we developed a three-dimensional organoid system based on our model. We generated uniform aggregates from day-12 TE cells and cultured them in suspension under agitation (Fig. 4f), inspired by a study showing that agitation promotes correct polarity in TB organoids, with STBs forming an outer layer around CTBs^54^. Organoids proliferated robustly in suspension and developed a clear internal organization. Staining for ITGA6, which broadly marks VCTs, and SDC1, which marks STBs, revealed discontinuous patches of SDC1+ syncytia encircling VCT domains, mirroring the layered pattern seen in the chorion (Figs. 4g and S6a).

Dual staining for VIM and GATA3 revealed that the majority of organoids comprised VIM+ cells typically concentrated toward the interior surrounded by layers of GATA3+ TB (Figs. 4h and S6b), reminiscent of the secondary villus topology of the early chorion. There was a variable VIM:GATA3 ratio in 3D cultures that could likely be further optimized by adjusting aggregation timing and size. We also found aggregates of VIM+ cells in day-18 adherent cultures, consistent with single cell profiling (Fig. S6b). Within the TB shell, HLA-G+ cells emerged spontaneously without directed induction of EVT maturation and localized to the outer perimeter of organoids, lining Ki67+ proliferative cells (Fig. 4i). Although we could not directly resolve distinct EVT populations in the single-cell transcriptomes, IF staining showed that organoids contained CDH5+ cells on the periphery, and adherent cultures also displayed ITGA1+ cells, which *in vivo* correspond to endovascular and interstitial EVT subtypes, respectively (Figs 4j and S6d). Finally, we tested whether the addition of A83-01 and neuregulin 1 (NRG1), components previously used for EVT maturation, affected organoid structure or organization. Neither polarity nor the syncytial or extravillous potential appeared different conditions (Fig. S6e).

Together, these findings demonstrate that HIPPO-driven differentiation culminates in the self-organization of chorionic structures that integrate both epithelial and stromal compartments. The resulting organoids reproduce hallmark features of the nascent chorionic villus, including spontaneous emergence of EVTs and internal mesenchymal core, establishing a platform for modeling early human placental architecture and function.

### Isolated columnar TBs demonstrate syncytial and extravillous potential *in vitro*

Lastly, we sought to demonstrate the power of our model to interrogate the emergence and properties of specific chorionic lineages. We focused on the villous column TB, a rare population at the base of anchoring villi bridging the epithelial and extravillous compartments. *In vivo*, these cells are thought to arise from VCT or VCTp populations, and are hypothesized to maintain a proliferative capacity with dual potential for both villous and invasive lineages^55^, although directly demonstrating their developmental potential has been elusive. We asked whether our system could capture this population and test its developmental capacity directly.

In the placenta, these cells are marked by *ITGA2*, however in our day-18 cultures, *ITGA2* was broadly expressed across several TB states, consistent with reports from stem-cell-derived TB organoids^56^. This difference in integrin switching between *in vivo* and *in vitro* possibly reflects a difference in the biophysical properties of the surrounding microenvironment, which we do not intentionally replicate. We therefore sought a marker that more selectively identifies the columnar TB population. Among ten most differentially expressed genes within the VCTc cluster, *CLDN6* was highly selective in our dataset and in the placenta reference to the columnar population (Fig. 5a). IF on day-18 adherent cultures confirmed that CLAUDIN-6 protein localizes to a discrete epithelial layer positioned above the ITGA6+ cells, reminiscent of the arrangement of columnar TB at anchoring villi (Fig. 5b). In chorionic organoids, we observed CLAUDIN-6 protein in the subset of cells both in the outer edge and the underlying layer, as we would expect for a transitioning VCT to EVT population (Figs 5c and S6d).

**Figure 5:**
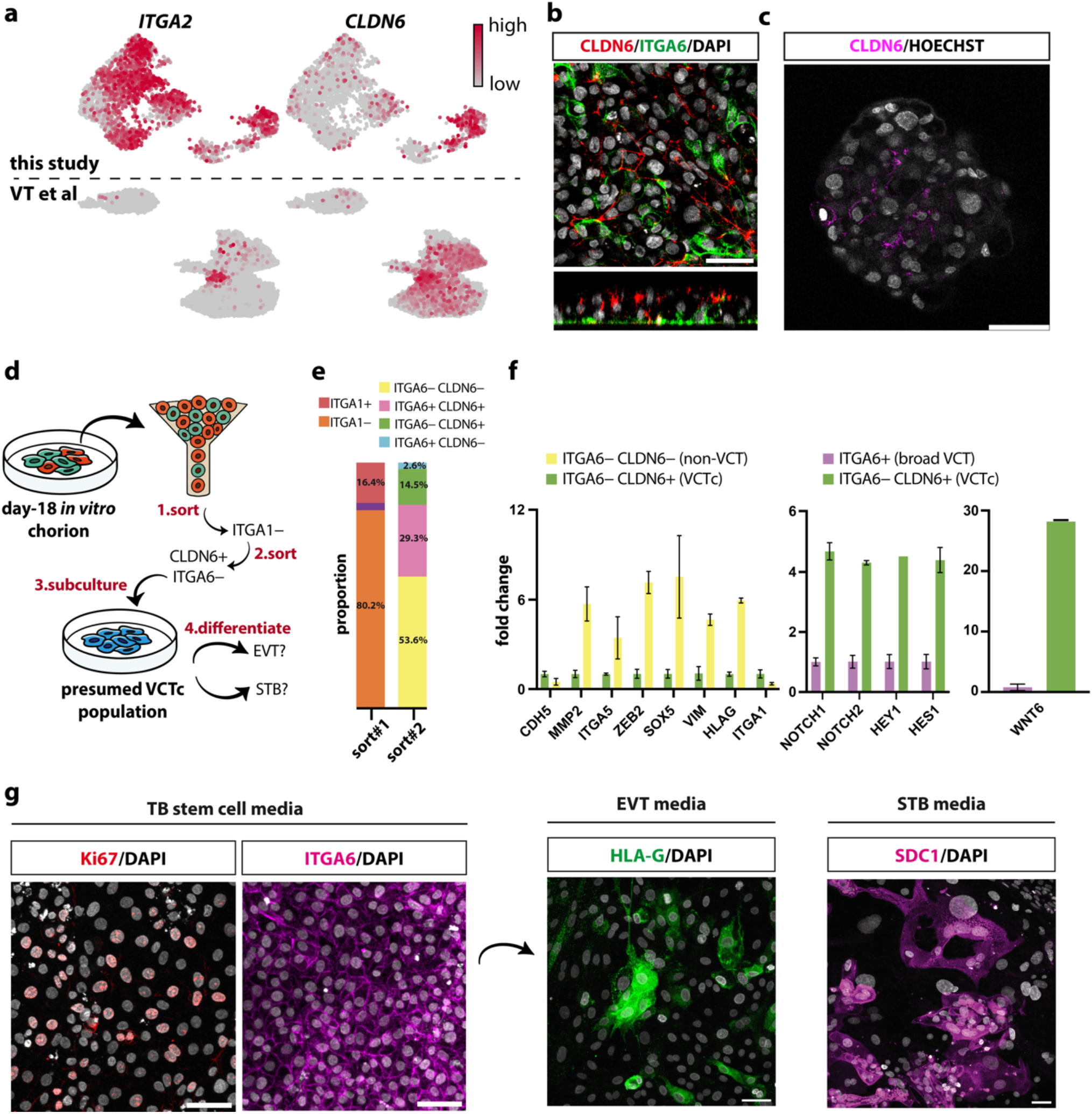
Isolation and lineage potential of CLDN6+ columnar TBs. (a) Expression of ITGA2 and CLDN6 in day 18 in vitro cultures compared with the first-trimester chorion reference^47^. (b-c) IF confirming discrete localization of CLAUDIN-6 above the ITGA6+ basal layer in day 18 cultures, as well as its localization in organoids. (d) Experimental workflow for the isolation and functional testing of columnar TBs. Day-18 adherent chorion cultures were sorted sequentially: first, ITGA1+ cells were removed to exclude differentiated EVTs; from the ITGA1− fraction, CLDN6+ITGA6− (columnar) subset was collected, expanded in TB stem cell medium, and subsequently differentiated under EVT or STB conditions. (e) Flow-cytometric quantification of sorted fractions (n > 2 replicates pooled per condition). (f) qPCR analysis showing relative expression of EVT (CDH5, MMP2, ITGA5, HLA-G), choriomesodermal (ZEB2, SOX5, VIM), and columnar (NOTCH1, NOTCH2, HEY1, HES1, WNT6) genes across various sub-populations (n = 2 biological replicates; bars show mean ± s.d.). (g) CLDN6+ITGA6− TBs cultured for two weeks in TB stem cell medium showing Ki67 and ITGA6 expression, followed by differentiation, showing HLA-G+ and SDC1+ expression. All scale bars, 50 µm.

To isolate this population, we designed a two-step sorting strategy to enrich bona fide columnar cells while excluding other chorionic lineages (Fig. 5d). We first removed ITGA1+ cells to deplete differentiated EVTs (Fig. 5e). From the ITGA1− fraction, we separated CLDN6+ITGA6−, CLDN6−ITGA6−, and ITGA6+ subsets (Fig. 5e). We considered the ITGA6+ cells to broadly represent VCT states, while CLDN6−ITGA6− represent the remaining populations: STB, EVT and chorionic mesoderm. Indeed, this population showed a significantly higher expression of EVT (*HLA-G, ITGA5*, and *MMP2*) and choriomesodermal (*SOX5, VIM, ZEB2*) genes, compared to the CLDN6+ITGA6− population (Fig. 5f).

Approximately 9% of cells were CLDN6+ ITGA6− ITGA1−; this fraction showed a characteristic column TB signature with elevated *NOTCH1, NOTCH2, HEY1, HES1* compared to the ITGA6+ villous subset, consistent with data from columnar cells sorted from the placenta^55^ and elevated NOTCH activity in the proximal column^57^ (Fig. 5f). We also checked for WNT6, which is selectively expressed in VCTc population in the first trimester placenta according to differential expression and, indeed, it was elevated in our sorted population (Fig. 5f). These data, together with single-cell profiling and IF, identify CLDN6+ ITGA6− ITGA1− cells as the columnar VCT population.

We then tested developmental potential. We subcultured CLDN6+ ITGA6− ITGA1− in TB stem cell medium^4^. After eight days, a fraction of cells retained Ki67+ proliferative capacity, however all cells regained ITGA6 expression, indicating a reacquisition of a VCTp phenotype (Fig. 5g). When exposed to lineage-specific cues, sorted cells efficiently and at high yield generated SDC1+ STB and HLA-G+ EVT lineages (Fig. 5g).

Together, these experiments directly demonstrate that CLDN6+ columnar TB possess both syncytial and extravillous differentiation potential, supporting their proposed^55^ bipotent identity within the chorionic lineage.

## Discussion

The molecular events that define the first lineage decision in the mammalian embryo remain incompletely understood. The initial trigger that separates the inner cell mass from the TE likely arises from early polarity cues or from preexisting asymmetries in RNA and protein distribution as early as the two-to four-cell stage, or from a combination of both^58^. Regardless of these early biases, the mechanistic effector that sets the TE trajectory in motion is the HIPPO-LATS-YAP axis. Here, we use targeted modulation of this pathway to show that its suppression is sufficient to initiate the human TE program and that the ensuing transcriptional sequence culminates in the formation of early chorionic lineages and quasi-architecture. In doing so, we sought to create an experimental framework that follows the developmental logic of the embryo rather than relying on initial BMP4 or WNT stimulation, which single-cell data of the early human embryo indicates is not active during TE induction. Because the framework starts from primed hESCs and transiently induces a window of TE competence, it offers an additional benefit of a highly standardized and widely accessible basis for reproducibly dissecting the molecular and mechanical principles that organize early placental development.

Curiously, both the integrated embryonic reference and our model show a comparable pattern of endogenous WNT activity downstream of HIPPO suppression. Noncanonical WNT ligands appear early in the TE trajectory, followed by the sequential activation of canonical ligands. Similarly, BMP ligands and direct targets of BMP signaling emerge later. These patterns suggest that WNT and BMP signaling are not required for TE induction but may act later to sustain the regenerative capacity of the chorionic villus. Consistent with this view, several studies have shown that WNT signaling supports the maintenance of TB stem cells^59,60^. The endogenous WNT activity may explain why ITGA6+ cells arise spontaneously in our cultures after TE specification, independent of exogenous WNT agonists, albeit to a lesser degree. The full extent of the roles of WNT and BMP in setting up the chorion is still unresolved. We speculate that BMP and WNT could help reinforce HIPPO pathway activity within the uterine environment, where increasing cell density and mechanical constraints are expected to attenuate nuclear YAP localization. Because these signals emerge endogenously downstream of HIPPO, our model provides an opportunity to probe the HIPPO-WNT-BMP interaction in experimental models of implantation.

In the human embryo, the secondary chorion forms when extraembryonic mesoderm lines the TB epithelium to generate villous projections. Although our model does not reproduce the morphogenetic steps of this process, the combined gene and protein expression, as well as computational evidence indicates that it reaches a comparable level of cellular complexity, containing the major TB and stromal populations of the early chorion. Although the TB fates appear to follow the developmental trajectory, the stromal lineage co-emerges with the TB. Multiple hypotheses exist on where chorionic mesoderm comes from, including from the primitive streak or from the hypoblast^61^, however its origin remains unresolved. A similar mesoderm-like population has been reported in TB stem cell cultures^62^, suggesting that the co-appearance of stromal and TB lineages may reflect an inherent plasticity of early extraembryonic tissue. Definitively resolving where chorionic mesoderm comes from in the human embryo would require lineage-tracing across peri-implantation and early post-implantation stages, e.g. using inducible CRISPR recorders introduced before implantation, followed by live imaging and spatial readouts through secondary villus formation. Such experiments would be highly challenging and they would require extended culture beyond the widely adopted ethical limits. Despite uncertain fidelity of chorionic mesodermal precursor in our model, we are excited that our model, especially in the 3D context, recreates the juxtaposition between the epithelial and stromal chorionic compartments offering a controlled setting to examine how their interactions initiate villus development.

Our model also allowed us to take a closer look at the columnar TB population. Columnar TBs at anchoring villi have been proposed as a proliferative precursor of EVTs with a potential to regenerate the villous CTB population^55^, although directly testing this hypothesis has been challenging. We identified a surface marker of this population, CLAUDIN-6, which allowed us to isolate it from living cultures and directly demonstrate they can revert to ITGA6+ cells with both STB and EVT potential. We hope our model and approach could set the stage to discover how the crosstalk between early chemical and mechanical signals bias fate at the column toward anchoring, invasion, or villous renewal, questions that link lineage control to implantation depth and early placental stability.

## Acknowledgments

We thank A. Danechi, S. Liu. E. Mattioli, and A. Misra for technical assistance, and the current and former members of the Simunovic lab for expertise and critical feedback. We thank A. Brivanlou (The Rockefeller University) for gifting the RUES1 and RUES2 lines. The study was supported by the Burroughs-Wellcome Fund Next Gen Pregnancy Initiative Award (NGP10152), The Blavatnik Fund for Engineering Innovations in Health Seed Fund, Allen Distinguished Investigator Award (PGAF 202211-13815), NYSCF Robertson Stem Cell Investigator Award (NYSCF-R-I80), and Columbia funds. We thank

B. Corneo, M. Kissner, and the staff of the Columbia Stem Cell Initiative Core for their expert assistance.

## Materials

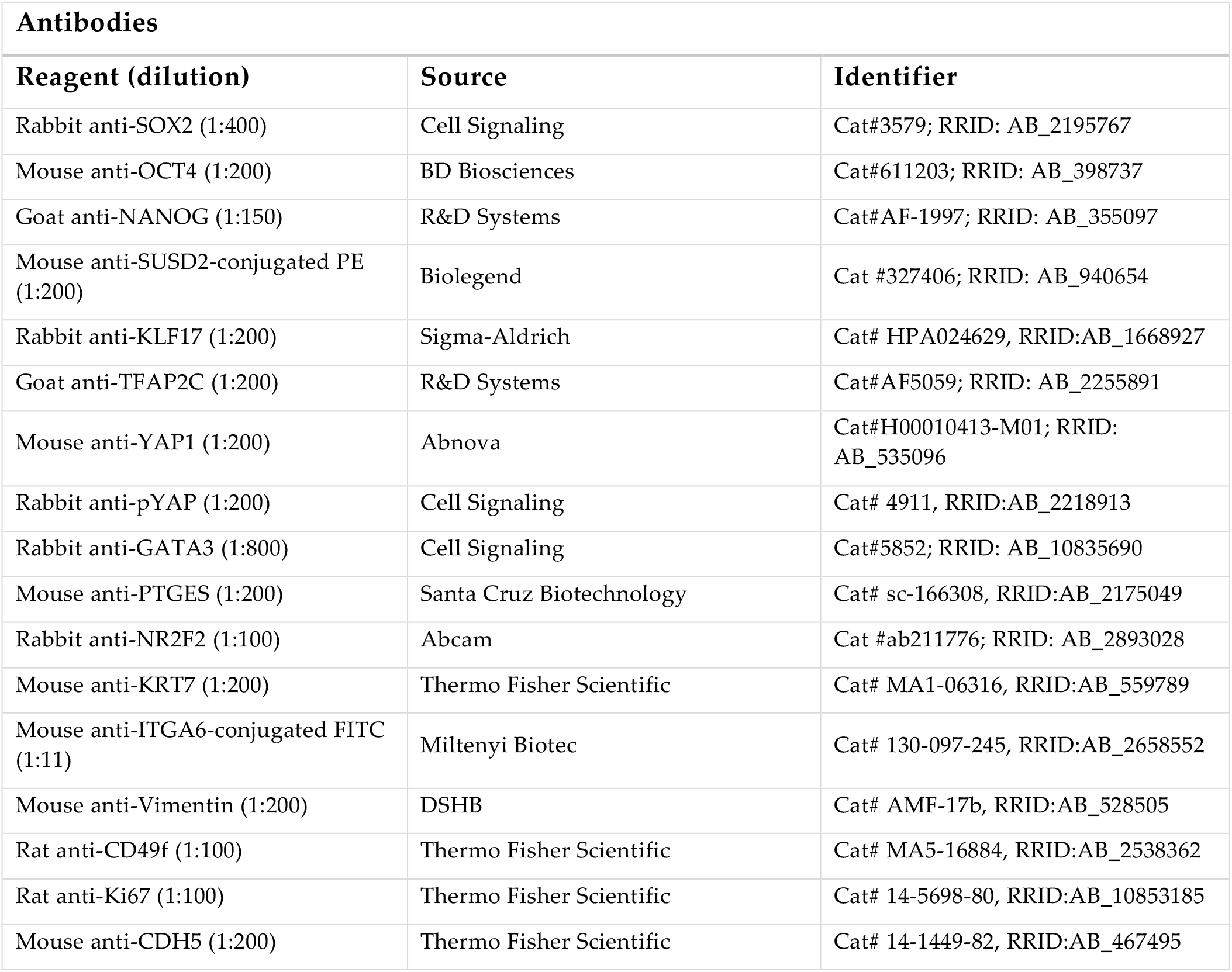
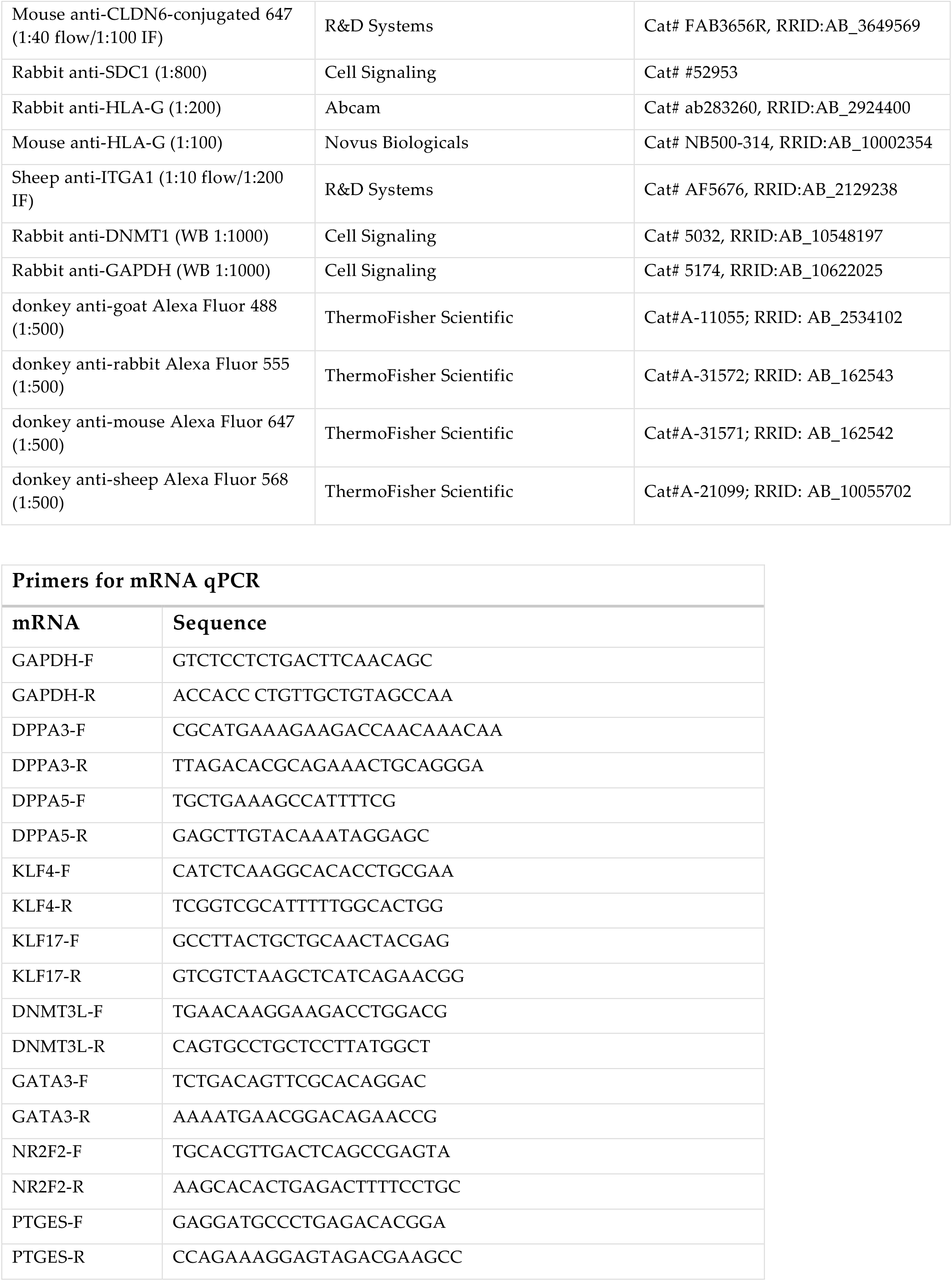
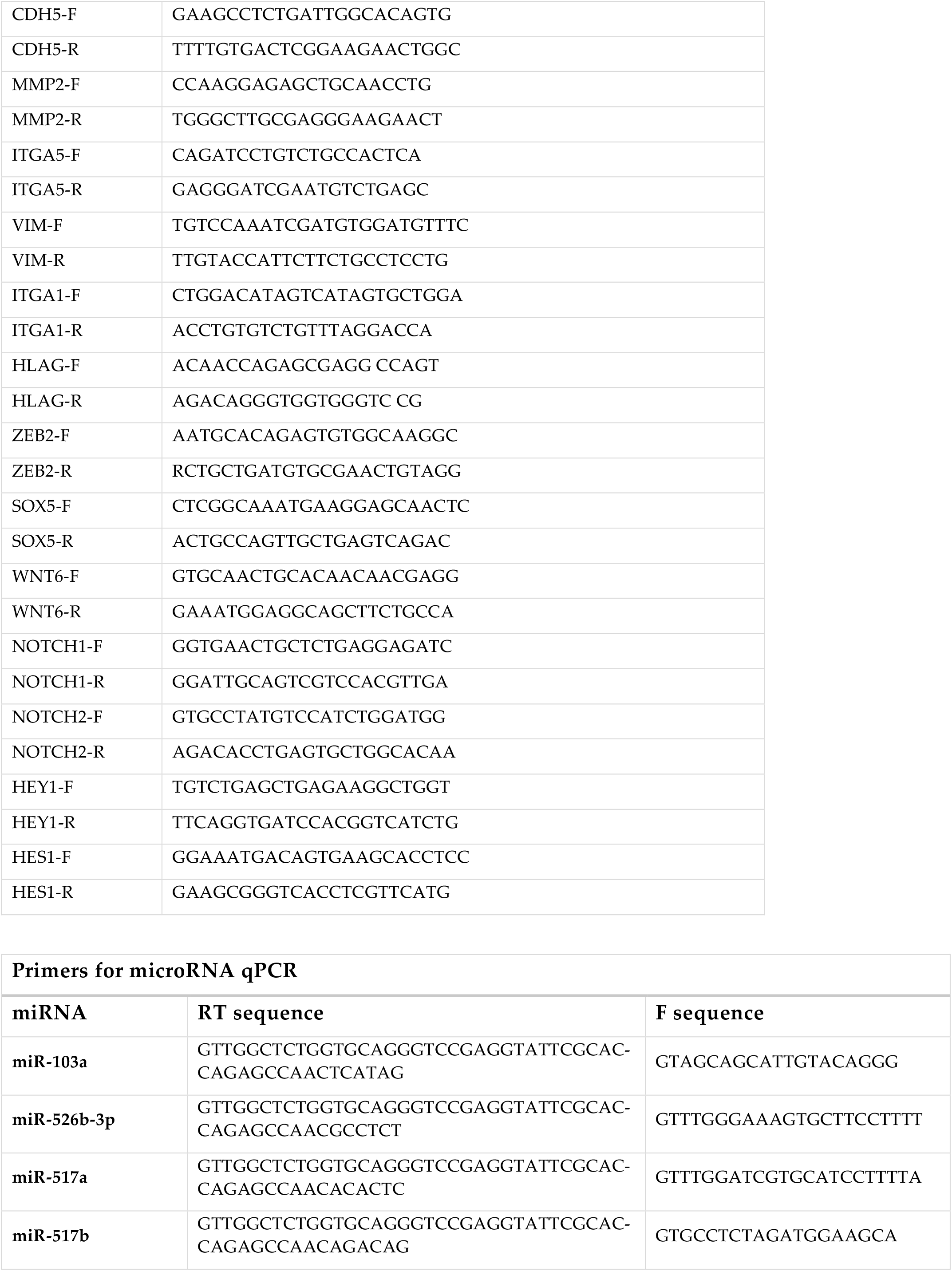
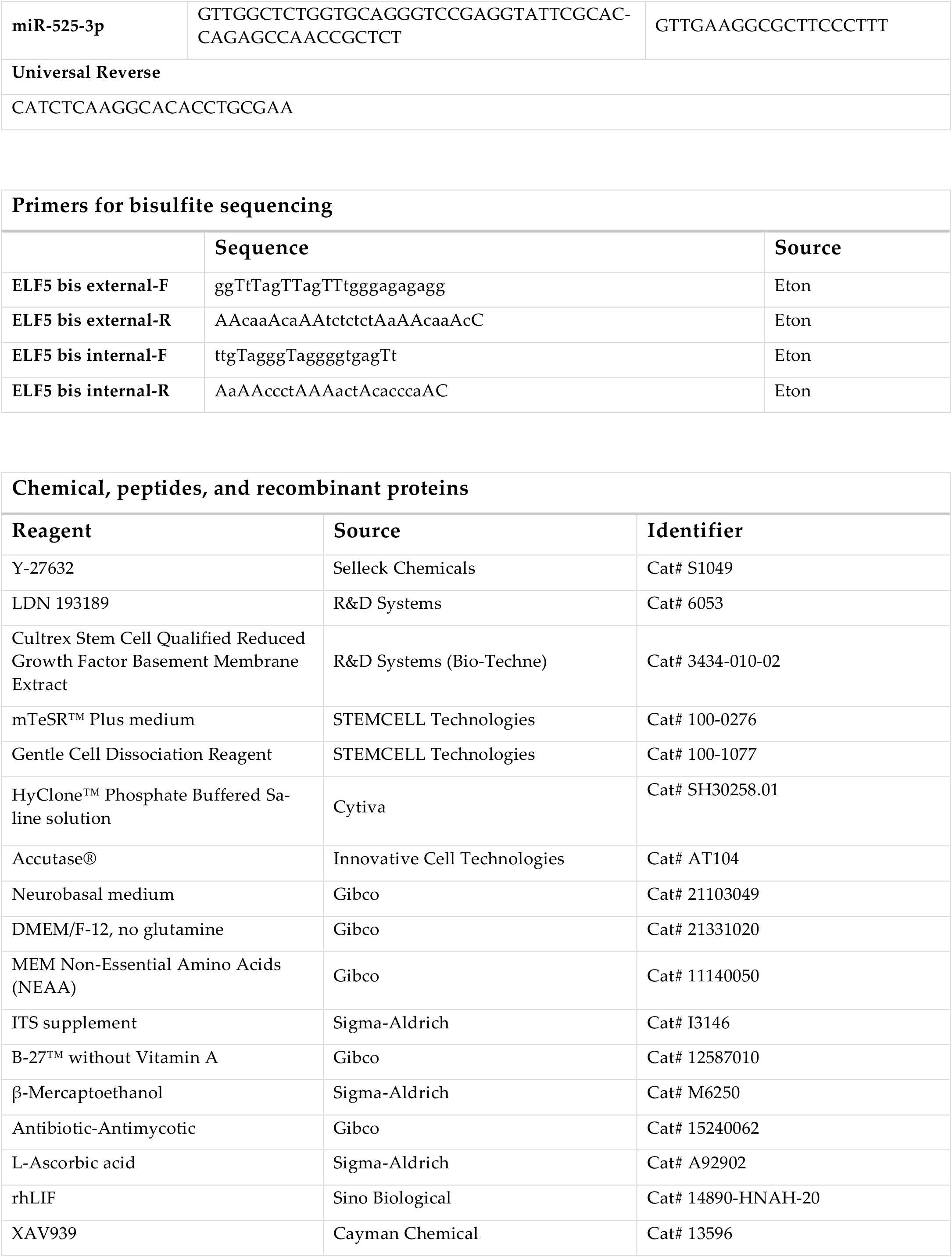
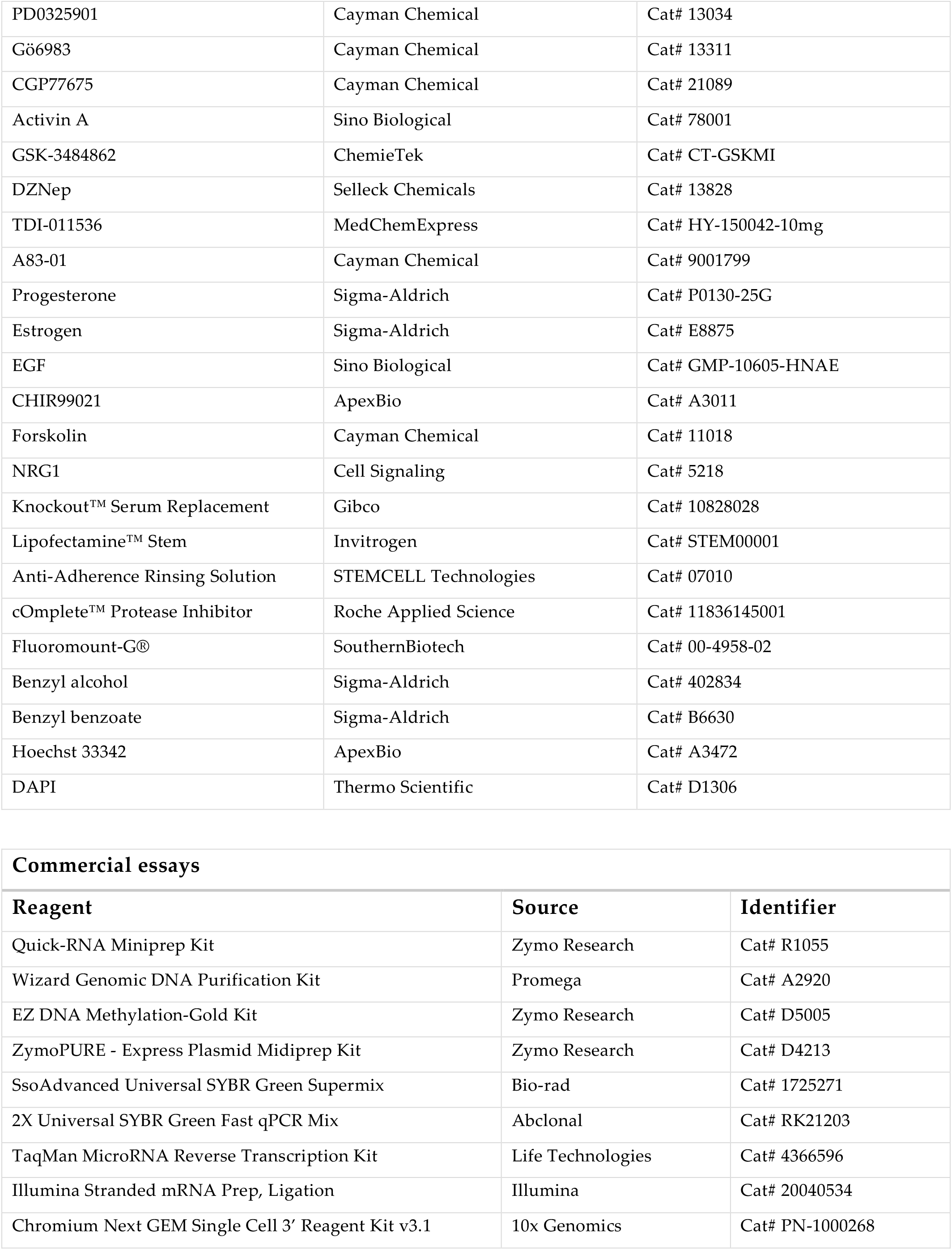
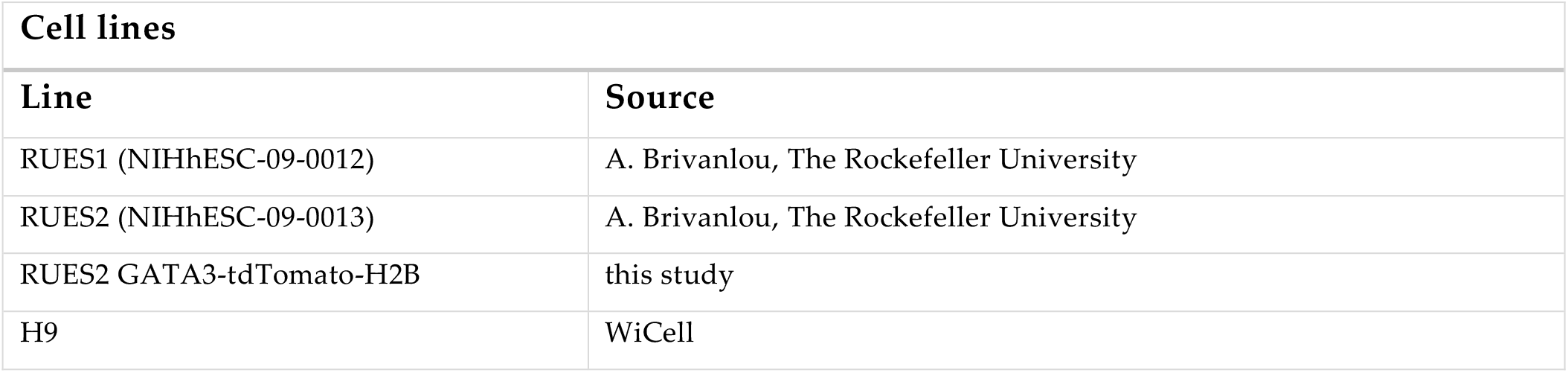

## Methods

### Ethics statement

The work reported in this paper underwent review by and received approval from the Columbia University Human Embryonic and Human Pluripotent Stem Cell Research Committee, which reviews protocols prospectively and on an ongoing basis. This paper does not use embryo models or natural embryos and only employs directed differentiation techniques, and is fully in line with the latest ISSCR guidelines on stem cell research.

### Cell culture

Four NIH-approved human embryonic stem cell lines were used in the study: RUES1, RUES2, H1 and H9. They were cultured under feeder-free conditions on Cultrex-coated tissue culture plates (1:100 dilution in DMEM/F12 (Cytiva)). Cells were maintained in mTeSR Plus medium (STEMCELL Technologies) and incubated at 37 °C in a humidified incubator with 5% CO_2_. Medium was changed daily. Cells were passaged every 3-4 days using Gentle Cell Dissociation Reagent (STEMCELL Technologies) following a PBS–/– (Cytiva) wash. 10 µM of ROCK inhibitor Y-27632 (Selleck Chemicals) was added during passaging and maintained for the first 24 hours post-split to promote survival. Cells were routinely screened for mycoplasma contamination and confirmed to be free of infection.

### Generation of RUES2-GATA3-Tdtomato reporter

GATA3-H2B-tdTomato-lox-PGK-puro-lox was constructed by Gibson assembly of five fragments (NEB-uilder HiFi DNA Assembly 2x Master Mix). Left and right homology arms were obtained by cloning 616 bp and 517 bp fragments, respectively, from hESC gDNA. The H2B-tdTomato fragment was cloned from pAAV-CAG-H2B tdTomato (Addgene plasmid #116870, a gift from Loren Looger). The PGK-puro fragment was cloned from pCW57.1-DUX4-WT (Addgene plasmid #99282, a gift from Stephen Tapscott). Lox sites were added in via primer amplification during cloning. The bacterial backbone fragment was cloned from pSpCas9(BB)-2A-GFP (PX458) (Addgene plasmid #48138, a gift from Feng Zhang). The plasmid assembly mix was transformed (NEB 5-alpha Competent E. coli high efficiency), and correct clones were purified (Zymochem Plasmid Midiprep kit, Plasmidsaurus whole plasmid sequencing). hESCs were transfected (Invitrogen Lipofectamine Stem), antibiotic-selected with puromycin (2 ug/mL), and single-cell cloned by seeding at low density (∼10 cells per cm^2^). Clonal hESC lines were stimulated with BMP4 to induce GATA3 expression to confirm proper integration of the fluorescent cassette at the GATA3 C-terminus.

### tHENSM+2i induction

To reset mTeSR-cultured hESC to a transient state with TE potential, they were plated as single cells onto Cultrex-coated 6-well plates at ∼30-50 × 10^3^ cells/cm^2^ in mTeSR medium supplemented with 10 µM Y-27632 and allowed to attach overnight. The following day, the medium was switched to HENSM supplemented with 2i and kept for 5-6 days, refreshing media daily (referred to as tHENS+2i). Composition, <u>basal medium</u>: 1:1 mixture of Neurobasal (Gibco):DMEM/F12 without HEPES and L-glutamine (Corning), supplemented with: 1× GlutaMAX (Gibco), 1× MEM non-essential amino acids (Gibco), 1× ITS (Sigma-Aldrich), 1× B27 without vitamin A (Gibco), 1× β-mercaptoethanol (Sigma-Aldrich), 1× Antibiotic-Antimycotic (Gibco), 50 µg/mL L-ascorbic acid (Sigma-Aldrich), and 0.2% (v/v) Cultrex (R&D Systems); <u>+ HENSM active ingredients (slightly optimized from published composition)</u>: 20 ng/mL rhLIF (Sino Biological), 2 µM XAV939 (Cayman Chemical Company), 1 µM PD0325901 (Cayman Chemical Company), 2 µM Gö6983 (Cayman Chemical Company), 1.2 µM CGP77675 (Cayman Chemical Company), 10 ng/mL Activin A (Sino Biological), and 1.2 µM Y-27632); <u>+ 2i</u>: 0.5 mM GSK-3484862 (DNMT1 inhibitor, ChemieTek) and 50nM DZNep (EZH2 inhibitor, Selleck Chemicals). Culture was carried out under 5% CO_2_ and 7% O_2_.

### TE induction

tHENSM+2i cells were passaged into single cells using Accutase (Innovative Cell Technologies) and seeded onto Cultrex-coated 6-well plates in TA medium with 10 µM Y-27632 added for the first 24 hours. TA medium composition, <u>basal medium</u>: 1:1 mixture of Neurobasal (Gibco):DMEM/F12 without HEPES and L-glutamine (Corning), supplemented with: 1× GlutaMAX (Gibco), 1× MEM non-essential amino acids (Gibco), 1× ITS (Sigma-Aldrich), 1× B27 without vitamin A (Gibco), 1× β-mercaptoethanol (Sigma-Aldrich), 1× Antibiotic-Antimycotic (Gibco), 50 µg/mL L-ascorbic acid (Sigma-Aldrich), and 0.2% (v/v) Cultrex (R&D Systems) <u>+ active ingredients</u>: 3 µM TDI-011536 (MedChem Express) ± 1 µM A83-01 (Cayman Chemical Company) for the first 6 days (see main text). Culture was carried out under 5% CO_2_ and 7% O_2_. Medium was replaced daily throughout the differentiation.

For perturbation experiments, XAV939 (2 µM) and LDN-193189 (1 µM) (R&D Systems) were added from TE induction and maintained continuously throughout the differentiation for a total of 12 days, with daily medium changes.

### Chorion induction

On day 12 of TE induction, cells were transitioned into <u>basal medium</u>: 1:1 mixture of Neurobasal (Gibco):DMEM/F12 without HEPES and L-glutamine (Corning), supplemented with: 1× GlutaMAX (Gibco), 1× MEM non-essential amino acids (Gibco), 1× ITS (Sigma-Aldrich), 1× B27 without vitamin A (Gibco), 1× β-mercaptoethanol (Sigma-Aldrich), 1× Antibiotic-Antimycotic (Gibco), 50 µg/mL L-ascorbic acid (Sigma-Aldrich), and 0.2% (v/v) Cultrex (R&D Systems), 1 µM Progesterone (Sigma-Aldrich), and 10 nM Estrogen (Sigma-Aldrich) ± 50 ng/ml EGF (Sino Biological) and 2 µM CHIR99021(ApexBio). Cells were kept in this medium for at least 6 additional days. Culture was carried out under 5% CO_2_ and 7% O_2_.

### Chorion organoids

On day 9 of TE induction (post tHENSM+2i), cells were dissociated with Accutase and seeded into AggreWell™ 400 plates (STEMCELL Technologies, pre-treated with Anti-Adherence Rinsing Solution) at a density of 300 cells per microwell in TE medium supplemented with 10 µM Y-27632. After cells settled to the bottom of the wells, the plate was centrifuged at 100×g for 3 minutes to enhance aggregation before incubation. On day 10, media was refreshed to TE medium without Y-27632. On day 12, aggregates were collected and maintained in 2 µM CHIR99021 + 50 ng/ml EGF, with media refreshed every other day until day 16, and then daily until day 20. On day 20, aggregates were transferred from AggreWell™ 400 plates to a 6-well tissue culture plate pre-treated with Anti-Adherence Rinsing Solution. Aggregates were maintained in suspension on an orbital shaker at low rpm for 48 hours. Cultures were maintained at 37 °C under 5% CO_2_ and 7% O_2_ throughout the procedure.

### Western blotting

Cells were washed with PBS -/- and harvested with Accutase. Cell lysates were prepared using Ripa buffer supplemented with 1× protease inhibitors (Roche Applied Science, #11836145001) and 0.1mM DTT. After centrifugation for 3 min at 16,000 g, the supernatant was boiled and equal amount of protein subjected to SDS-PAGE, electro-transferred onto PVDF membrane and visualized by ECL (GE Healthcare). Immunodetection was performed using the following primary antibodies: anti-DNMT1 (Cell Signaling #5032) and GAPDH (Cell Signaling #5174). Chemiluminescence was captured by the ChemiDoc MP imager (BioRad).

### Quantitative PCR for mRNA

Total RNA was extracted using the Quick-RNA Miniprep Kit (Zymo Research) following the manufacturer’s protocol. First-strand cDNA was synthesized from 100 ng of RNA using All-In-One 5× RT MasterMix with gDNA removal (ABM) according to the manufacturer’s instructions. The resulting cDNA was diluted 10-fold in nuclease-free water. Real-time qPCR was performed on a Bio-Rad CFX96 Real-Time PCR Detection System using either SsoAdvanced Universal SYBR Green Supermix (Bio-Rad) or 2X Universal SYBR Green Fast qPCR Mix (Abclonal), following the respective protocols. Samples were run in technical replicates. Gene expression levels were normalized to GAPDH as an endogenous control and reported as ΔΔCq.

### Quantitative PCR for microRNA

Quantification of C19MC miRNAs was performed by adapting a published method^63^. Briefly, 10 ng of total RNA was reverse transcribed using miRNA-specific stem-loop RT primers obtained from the primer sets as previously described^41^, using the TaqMan MicroRNA Reverse Transcription Kit (Life Technologies) according to the manufacturer’s instructions. The resulting cDNA was quantified by real-time PCR using SYBR Green chemistry as described above. All reactions were run in technical replicates. Expression levels were normalized to hsa-miR-103a^64^ and reported as ΔΔCq.

### Bisulfite Sequencing

DNA was isolated from cells using the Wizard Genomic DNA Purification Kit (Promega). 400ng of DNA was then converted by bisulfite using the EZ DNA Methylation-Gold kit (Zymo). [−136 - +238] region of *ELF5* promoter was amplified by sequential PCR using nested primers^10^ and the PCR product was ligated into the pGEM-T vector (Promega). Eight clones were sequenced for each condition. Sequencing results were analyzed and visualized using QUMA^65^.

### Immunocytochemistry

The list of primary and secondary antibodies, antibody dilutions, and antibody catalogue numbers can be found in Table S1. IF staining was performed following standard protocols. Briefly, adherent colonies or organoids were fixed in 4% paraformaldehyde (in PBS+/+) for 30 min at room temperature, then permeabilized and blocked for 1h in 2% BSA and 0.1% Triton X-100 prepared in PBS+/+. Primary antibodies were diluted in blocking buffer and incubated with samples overnight at 4°C. The following day, samples were washed three times with 0.1% Triton X-100 in PBS+/+, then incubated for 1h at room temperature in the dark with fluorophore-conjugated secondary antibodies and DAPI (Thermo Scentific) or Hoechst 33342 (ApexBio), also diluted in blocking buffer. Coverslips were washed again and mounted using Fluoromount-G (SouthernBiotech), then sealed with nail polish. For 3D organoids, staining was carried out in suspension in 1.5 mL Eppendorf tubes using the same protocol. After staining, organoids were transferred to 8-well Ibidi imaging chambers.

### Clearing of 3D organoids

Prior to clearing, organoids were washed in distilled water and dehydrated through a graded methanol series (25%, 50%, 75%, 100%, v/v in water), with 5-10 minutes per step. Samples were optically cleared using a 1:2 mixture of benzyl alcohol and benzyl benzoate (BABB, Sigma-Aldrich) at room temperature until transparent and then they were immediately imaged.

### Microscopy imaging

Epifluorescence and confocal microscopy imaging was performed using a Leica Stellaris microscope with either a 20× dry or 25× water-immersion objective. Image processing and rendering were performed using Fiji (ImageJ) or Leica LAS X. Live-cell imaging was done under controlled environmental conditions using an Oko labs tri-gas cell incubator (37°C, 5% CO_2_, normoxia) using a 20× dry objective.

### Flow Cytometry

The list of primary and secondary antibodies, antibody dilutions, and antibody catalogue numbers can be found in Table S1. Cells were digested into single cells using Accutase and washed with FACS buffer (2% FBS and 10 µM Y-27632 in DMEM/F12). Cells were resuspended in FACS buffer at appropriate density, and primary antibodies were added for 1h on ice. After washing, cells were incubated with secondary antibodies and Hoechst 33342 (Apex Bio) for 30 minutes on ice. The samples were washed, filtered through 40 µm cell strainers, and analyzed using an Agilent NovoCyte Penteon or sorted using a Sony MA900 equipped with a 100 µm nozzle. Data were processed using FlowJo software.

### Bulk RNA-sequencing and data analysis

Cells were collected and lysed under various conditions (mTeSR, tHENSM+2i, tHENSM, then 3d, 6d, 9d, and 12d of TE induction), with *n* = 3 for all except *n* = 2 for day 6. Total RNA was extracted at defined time points during differentiation using the Quick-RNA Miniprep Kit (Zymo Research) following the manufacturer’s protocol. RNA purity and quantity was evaluated using a NanoDrop 2000 spectrophotometer (Thermo Fisher Scientific). Approximately 500 ng of total RNA per sample was used for library preparation with the Illumina Stranded mRNA Prep Kit (Illumina), according to the manufacturer’s instructions. Sequencing was performed on an Illumina NovaSeq X Plus system with paired-end 150 bp reads (PE150). Raw reads were quality filtered and trimmed using fastp and aligned to the human reference genome (hg38) using HISAT2 v2.0.5. Gene-level expression was quantified with featureCounts v1.5.0-p3, and FPKM values were calculated for normalization. Differential expression analysis was performed using DESeq2 v1.20.0 for samples with replicates. Genes with adjusted P ≤ 0.05 and |log_2_FC| ≥ 1 were considered significant. Low-expression genes were filtered out prior to analysis. Heatmaps, PCA, correlation, and clustering analyses were performed using R v3.5.1

To generate PCA plot comparing tHENSM+2i with naïve and primed-cultured hPSCs, we curated publicly available bulk RNA-seq datasets, representing various pluripotent stem cell states^9,66–68^: GSE153212, GSE138304, GSE150772, CNP0001454. We removed samples representing repeated cell lines under identical culture conditions to avoid redundancy. We filtered out genes with zero variance, low average expression, as well as mitochondrial, ribosomal, pseudogene, heat shock, and housekeeping genes in PCA analysis. We used ComBat (*sva* package) to apply batch correction, with study of origin specified as the batch variable. Finally, we calculated PCs on 10,000 most variable genes, scaled by Z-score.

We used the GSEA function in the *clusterProfiler69* R package^69^ to perform enrichment analysis on differentially expressed genes from day-12 vs day-0 samples, using a curated list of YAP/TAZ targets. Enrichment scores and *p*-value was reported, and enrichment curves were generated with *enrichplot*.

To quantify transposable elements, we followed a published approach to detect repeat families in bulk RNA-seq data^70^. In short, we constructed ‘repeatomes’ from *RepeatMasker* genome-wide annotation files for assembly hg38. We aligned bulk RNA reads to repeat genome sequences using *Bowtie2* (default parameters), with any alignment to a repeat family being scored.

To assess whether tHENSM+2i cells preserve imprinting, we performed allele-specific expression (ASE) analysis on bulk RNA-seq from primed and tHENSM+2i hPSCs. BAM files were preprocessed with the Genome Analysis Toolkit (GATK v4.3.0.0), including read group assignment (*AddOrReplaceReadGroups*) and splicing-aware correction (*SplitNCigarReads*). Variants were called per sample using *HaplotypeCaller* in RNA-seq mode (EMIT_VARIANTS_ONLY) with a minimum confidence threshold of 30. To restrict the analysis to loci informative for imprinting, we first identified heterozygous variants using *bcftools* and intersected these with a curated set of imprinted genes. Multiallelic and duplicate records were decomposed and normalized (*bcftools* norm), and spanning deletions and non-SNP variants were excluded. ASE counts were obtained with *ASEReadCounter*, yielding reference and alternate allele read counts at each heterozygous position. For downstream analysis, variant-level counts were aggregated at the gene level. Within each gene, the major allele was consistently assigned as reference to ensure orientation consistency, and a reference allele ratio compared to total was then computed, yielding 0.5 (biallelic) to 1.0 (monoallelic). Replicates were combined per condition, and gene-level ratios were summarized as medians.

### Single cell RNA sequencing and data processing

Cells for single cell RNA profiles were harvested on days 6, 12, and 18 of treatment, as depicted in Fig. 3b. For each time point, at least three biological replicates were pooled. At each time point, cultures were washed with PBS, incubated with Accutase at 37 °C for 5-10 minutes, and gently triturated to obtain a single-cell suspension. Dissociation was monitored microscopically to ensure minimal clumping, and filtered suspensions through a 40 µm strainer to remove residual aggregates. Cells were manually counted using TrypanBlue to dye the cells. Single-cell RNA-sequencing libraries were generated using the Chromium Next GEM Single Cell 3’ Reagent Kit v3.1 (10x Genomics) according to the manufacturer’s instructions. Library quality and fragment size were assessed using an Agilent 2100 Bioanalyzer and Qubit fluorometer (Thermo Fisher). Equimolar libraries were pooled and sequenced on an Illumina NovaSeq 6000 platform to generate paired-end 150 bp reads at an average depth of ∼20,000 reads per cell. Cell Ranger Count v8.0.1 (10x Genomics) was used to align reads to the human reference genome GRCh38 using default parameters.

For each dataset, droplets were evaluated with valiDrops (v1.0)^71^, which assigns quality-control status and live/dead labels to each barcode. To mitigate ambient RNA contamination, we used SoupX (v1.6)^72^. Each dataset was clustered in Seurat using standard preprocessing (log-normalization, variable feature selection, scaling, PCA, and Louvain clustering), and cluster assignments were transferred to the SoupX objects. Contamination fractions were estimated using curated sets of ribosomal (RPL, RPS), collagen (COL1A1, COL1A2, COL3A1), and actin (ACTB, ACTG1) genes that were detected but not expected to be endogenously expressed in all clusters. Valid gene sets were identified with *estimateNonExpressingCells*, and cell-specific contamination fractions were calculated with *calculateContaminationFraction*. Corrected count matrices were produced using *adjustCounts* and re-imported into Seurat for downstream analysis. We then quantified multiple QC features for each cell, including total detected genes (nFeature_RNA), UMI counts (nCount_RNA), mitochondrial transcript percentage ribosomal transcript percentage, transcript complexity (log10GenesPerUMI), and a Gini coefficient of expression distribution. Thresholds for filtering were defined adaptively using ±3 median absolute deviations (MAD) from the dataset median for each metric, with upper cutoffs applied to mitochondrial and Gini values and lower cutoffs applied to complexity. Cells outside these thresholds were removed. Doublets were identified with scDblFinder (v1.12)^73^ applied to SingleCellExperiment objects converted from Seurat.

Final filtered datasets were log-normalized, top 2000 highly variable features were selected using the variance-stabilizing transformation, and data were scaled while regressing out sequencing depth and mitochondrial percentage. Dimensionality reduction was performed with PCA, and neighborhood graphs were constructed using the first 50 PCs (FindNeighbors). Cells were clustered using the Louvain algorithm at resolution = 0.5 (FindClusters), and low-dimensional embeddings were generated with UMAP.

### Projection onto the six-dataset embryonic reference map

The embryonic reference map, constructed from six published single-cell datasets of the early human embryo, and its associated projection pipeline have been described previously and released as an opensource resource^20^. In Fig. 3b we replotted the UMAP representation of the six-dataset embryonic reference, using the original annotations. After stringent QC of our three *in-vitro* single-cell transcriptomes, we projected those datasets onto the reference using the published projection tool, with parameter modifications to suit our data. For Milo aggregation, we used the first 50 PCs with graph_k = 15, graph_d = 40, nhood_k = 10, nhood_d = 40, prop in the range 0.30-0.75, depending on dataset, refined = TRUE, and a fixed seed (456). We chose parameters to preserve finer-grained overlapping neighborhoods (∼700 per dataset) and to avoid clumping similar cells into overly broad aggregates.

The remaining steps followed the published pipeline: (i) alignment of query genes to the reference gene universe; (ii) deconvolution-based size-factor normalization and log-transformation without recentering (center.size.factors = FALSE), followed by cosine normalization on the reference high variable gene (HVG) set; (iii) projection of the query into the reference SVD basis (50 PCs); (iv) scoring of candidate anchors by Spearman correlation in the HVG space, with retention of anchors above a correlation threshold (we used 0.60-0.65 to prioritize high-confidence matches); (v) correction of the query embeddings by mutual nearest neighbor (MNN) pairing (*k* = 20-30, tricube kernel with ndist = 3), with average correction vectors estimated via a Gaussian kernel (*σ* = 0.1) and shift-variance adjustment, after which the corrected query PCs were concatenated with the reference fastMNN space; and (vi) embedding of the corrected query using the saved UMAP transform from the reference (seed = 10), with axes swapped to match the orientation to the published map.

### Trajectory and dendrogram inference from the embryonic reference map

To reconstruct lineage trajectories, we first excluded cell types not relevant to our analysis (excluded: all cell types prior to morula and the entire CS7 dataset apart from the amnion and primitive streak clusters), and cells with ambiguous signatures. We generated a force-directed layout to visualize global developmental trajectories. Diffusion maps were computed from MNN-corrected embeddings using Palantir (v1.2) with 55 components, *k* = 30 nearest neighbors, and seed = 12. A multiscale diffusion space was derived from the top 16 eigenvectors. The neighborhood graph was reconstructed in this space with Scanpy (v1.10), and the ForceAtlas2 algorithm was applied for 3,200 iterations with gravity = 1.4 and seed = 12. The resulting two-dimensional coordinates were used for visualization. We used scFates (v1.0)^43^ to generate a lineage dendrogram by first fitting a principal tree to the multiscale diffusion space derived from Palantir. We applied the SimplePPT algorithm using 190 nodes with regularization parameters *σ* = 0.20 and *λ* = 2.8, with the precomputed k-nearest neighbor graph built on the Palantir embedding. The learned principal tree was projected onto the ForceAtlas2 layout for visualization, from which the root node (morula) was inferred and specified to compute pseudotime. Short spurious branches (<3 nodes) were pruned, and a dendrogram representation was then generated to summarize branching structure and lineage relationships using the scFates.tl.dendrogram function. We assessed the association of gene expression dynamics with pseudotime using scFates.tl.test_association, enabling identification of genes with non-random pseudotemporal structure. We applied scFates.tl.fit to fit smoothed expression trends for all genes across trajectory segments. These fitted trends were visualized overlaid on dendrograms.

### Gene ordering along the pseudotime trajectory and visualization

To quantify temporal dynamics of differentiation, we subset the embryonic map to the trophoblast branch (morula, prelineage, TE, CTB, STB, ETV) and used fitted gene expression values derived from the scFates smoothed dendrogram representation along the trophoblast trajectory. For each gene, we smoothened fitted expression profiles across pseudotime bins using Gaussian filtering. Two statistics were combined: the pseudotime location of the expression peak (τ_peak) and the weighted median of the distribution (τ_med). A convex combination of these (*τ* = 0.5·τ_peak + 0.5·τ_med) provided a robust pseudotime index. We then sorted genes by τ to establish a continuous temporal order (earliest → latest).

To split the trajectory into early, transit and late genes, we computed the mean fitted expression profile across all genes and its derivative with respect to pseudotime. The transit region was defined as the region of maximum slope. The start and end of the transit period was defined where derivative dropped below 50% of its maximum, and the center was taken as the midpoint of this interval. We divided the τ-ordered gene list into three blocks of equal size: early genes: the first 10% of the ordered list, middle genes: 10% of transit genes equidistantly distributed around the detected transition, late genes: the first 10% of the late. For each block, fitted gene expression values were z-scored per gene (row-wise) and averaged across all genes in the block to yield a mean trajectory. Standard error of the mean (SEM) was computed at each pseudotime bin. The block trajectories were plotted with mean ± SEM shading.

The in vivo τ-ordered subsets (early/mid/late) were used to subset bulk RNA-seq data from differentiation time courses (d0, d3, d6, d9, and d12). Counts were variance-stabilized (DESeq2 VST) and averaged across replicates. For visualization, cubic splines were fit to expression values across timepoints, interpolated to 60 evenly spaced pseudotime bins, and row-wise z-scored. Heatmaps were generated to display early/mid/late subsets in direct correspondence with the *in vivo* ordering.

### Extending the embryonic reference map with first trimester placenta transcriptome

To extend the original embryonic reference, we included the Vento-Tormo dataset of the first trimester placenta (VT-FTP)^47^. We first subset the dataset to trophoblast and fibroblast lineages (as originally annotated) and downsampled the dominant VCT cluster to 4,000 cells to avoid over-representation. We then performed a single round of Milo neighborhood aggregation: we selected 4,000 highly variable genes, ran PCA with 50 components, built neighborhoods (*k* = 20, *d* = 50), and aggregated raw counts across neighborhoods to generate pseudocells. We normalized pseudocells by deconvolution size factors and log-transformed them, assigning each pseudocell a majority-vote cell type label. We harmonized the VT-FTP pseudocells to the gene universe of the six-dataset reference by zero-filling missing genes and reordering to match the reference. We rescaled all seven datasets with *multiBatchNorm*, preserving size factors and bringing library sizes to the scale of the lowest-coverage batch.

We defined integration features by extending the original 4,000 HVGs from the embryo reference. We added ∼1000 HVGs identified within VT pseudocells and supplemented them with ∼300 HVGs identified by differential expression between amnion (CS7) and VCT (VT). This produced a master set of ∼5,300 genes. We integrated all seven datasets with the published manual fastMNN pipeline^47^. In brief, we applied cosine normalization on the HVG set, centered each batch on the grand mean, scaled by the square root of the number of cells, performed global SVD (50 PCs), and merged batches sequentially in developmental order (Yan → Meistermann → Petropoulos → Guo → Xiang → Tyser → Vento-Tormo; *k* = 55, tricube weighting, *n*_dist_ = 3). We stored correction vectors, grand means, the SVD basis, and batch offsets for downstream projection.

We computed a UMAP on the corrected PC space and retained the transform model for query mapping. To improve separation of amnion and VCT while maintaining global developmental continuity, we systematically tuned UMAP parameters and selected n_neighbors = 85–105 and min_dist = 0.30–0.35.

### Projection of query datasets onto the extended reference map

We projected four query datasets onto the extended reference: two embryonic control datasets to demonstrate the fidelity of the reference, and the day-12 and day-18 *in vitro* transcriptomes. The control datasets were: (i) our previously published E10 human embryo single-cell RNA-seq^52^, and (ii) the single-nucleus dataset labeled ashley_collection_sn in the placenta atlas^53^, neither of which were used to construct the embryonic reference map.

To prepare each query dataset, we first harmonized the gene set to match the reference gene universe. We then normalized counts by deconvolution (using *scran*::computeSumFactors), applied log transformation without recentering and cosine-normalized expression values on the extended HVG set (∼5,300 genes). We then carried out the projection using the published Zhao *et al*. pipeline^20^, as above, with a Spearman correlation cutoff of 0.5.

### Use of generative AI

ChatGPT (versions 3, 4, and 5, including their sub-versions; OpenAI) was used to assist with troubleshooting code syntax and scripting during data analysis, and to refine hand-drawn cartoons in figures.

## Bibliography

1. Rossant, J., and Tam, P.P.L. (2022). Early human embryonic development: Blastocyst formation to gastrulation. Dev. Cell 57, 152–165. 10.1016/j.devcel.2021.12.022.

2. Infertility Prevalence Estimates, 1990-2021 (2023). 1st ed. (World Health Organization).

3. World Health Organization (2019). Trends in maternal mortality 2000 to 2017: estimates by WHO, UNICEF, UNFPA, World Bank Group and the United Nations Population Division (World Health Organization).

4. Okae, H., Toh, H., Sato, T., Hiura, H., Takahashi, S., Shirane, K., Kabayama, Y., Suyama, M., Sasaki, H., and Arima, T. (2018). Derivation of Human Trophoblast Stem Cells. Cell Stem Cell 22, 50–63 e6. 10.1016/j.stem.2017.11.004.

5. Xu, R.H., Chen, X., Li, D.S., Li, R., Addicks, G.C., Glennon, C., Zwaka, T.P., and Thomson, J.A. (2002). BMP4 initiates human embryonic stem cell differentiation to trophoblast. Nat Biotechnol 20, 1261–1264. 10.1038/nbt761.

6. Amita, M., Adachi, K., Alexenko, A.P., Sinha, S., Schust, D.J., Schulz, L.C., Roberts, R.M., and Ezashi, T. (2013). Complete and unidirectional conversion of human embryonic stem cells to trophoblast by BMP4. Proc Natl Acad Sci U A 110, E1212–21. 10.1073/pnas.1303094110.

7. Soncin, F., Morey, R., Bui, T., Requena, D.F., Cheung, V.C., Kallol, S., Kittle, R., Jackson, M.G., Farah, O., Dumdie, J., et al. (2022). Derivation of functional trophoblast stem cells from primed human pluripotent stem cells. Stem Cell Rep. 17, 1303–1317. 10.1016/j.stemcr.2022.04.013.

8. Seetharam, A.S., Vu, H.T.H., Choi, S., Khan, T., Sheridan, M.A., Ezashi, T., Roberts, R.M., and Tuteja, G. (2022). The product of BMP-directed differentiation protocols for human primed pluripotent stem cells is placental trophoblast and not amnion. Stem Cell Rep. 17, 1289–1302. 10.1016/j.stemcr.2022.04.014.

9. Bayerl, J., Ayyash, M., Shani, T., Manor, Y.S., Gafni, O., Massarwa, R., Kalma, Y., Aguilera-Castrejon, A., Zerbib, M., Amir, H., et al. (2021). Principles of signaling pathway modulation for enhancing human naive pluripotency induction. Cell Stem Cell 28, 1549–1565 e12. 10.1016/j.stem.2021.04.001.

10. Viukov, S., Shani, T., Bayerl, J., Aguilera-Castrejon, A., Oldak, B., Sheban, D., Tarazi, S., Stelzer, Y., Hanna, J.H., and Novershtern, N. (2022). Human primed and naïve PSCs are both able to differentiate into trophoblast stem cells. Stem Cell Rep. 17, 2484–2500.

11. Dong, C., Beltcheva, M., Gontarz, P., Zhang, B., Popli, P., Fischer, L.A., Khan, S.A., Park, K.M., Yoon, E.J., Xing, X., et al. (2020). Derivation of trophoblast stem cells from naive human pluripotent stem cells. Elife 9. 10.7554/eLife.52504.

12. Jang, Y.J., Kim, M., Lee, B.-K., and Kim, J. (2021). Induction of human trophoblast stem-like cells from primed pluripotent stem cells. bioRxiv.

13. Horii, M., Li, Y., Wakeland, A.K., Pizzo, D.P., Nelson, K.K., Sabatini, K., Laurent, L.C., Liu, Y., and Parast, M.M. (2016). Human pluripotent stem cells as a model of trophoblast differentiation in both normal development and disease. Proc. Natl. Acad. Sci. 113, E3882–E3891. 10.1073/pnas.1604747113.

14. Li, Y., Moretto-Zita, M., Soncin, F., Wakeland, A., Wolfe, L., Leon-Garcia, S., Pandian, R., Pizzo, D., Cui, L., Nazor, K., et al. (2013). BMP4-directed trophoblast differentiation of human embryonic stem cells is mediated through a DeltaNp63+ cytotrophoblast stem cell state. Development 140, 3965–3976. 10.1242/dev.092155.

15. Wei, Y., Wang, T., Ma, L., Zhang, Y., Zhao, Y., Lye, K., Xiao, L., Chen, C., Wang, Z., Ma, Y., et al. (2021). Efficient derivation of human trophoblast stem cells from primed pluripotent stem cells. Sci Adv 7. 10.1126/sciadv.abf4416.

16. Guo, G., Stirparo, G.G., Strawbridge, S.E., Spindlow, D., Yang, J., Clarke, J., Dattani, A., Yanagida, A., Li, M.A., Myers, S., et al. (2021). Human naive epiblast cells possess unrestricted lineage potential. Cell Stem Cell 28, 1040–1056 e6. 10.1016/j.stem.2021.02.025.

17. Cinkornpumin, J.K., Kwon, S.Y., Guo, Y., Hossain, I., Sirois, J., Russett, C.S., Tseng, H.W., Okae, H., Arima, T., Duchaine, T.F., et al. (2020). Naive Human Embryonic Stem Cells Can Give Rise to Cells with a Trophoblast-like Transcriptome and Methylome. Stem Cell Rep. 15, 198–213. 10.1016/j.stemcr.2020.06.003.

18. Osnato, A., Brown, S., Krueger, C., Andrews, S., Collier, A.J., Nakanoh, S., Quiroga Londoño, M., Wesley, B.T., Muraro, D., Brumm, A.S., et al. (2021). TGFβ signalling is required to maintain pluripotency of human naïve pluripotent stem cells. eLife 10, e67259. 10.7554/eLife.67259.

19. Io, S., Kabata, M., Iemura, Y., Semi, K., Morone, N., Minagawa, A., Wang, B., Okamoto, I., Nakamura, T., Kojima, Y., et al. (2021). Capturing human trophoblast development with naive pluripotent stem cells in vitro. Cell Stem Cell 28, 1023–1039 e13. 10.1016/j.stem.2021.03.013.

20. Zhao, C., Plaza Reyes, A., Schell, J.P., Weltner, J., Ortega, N.M., Zheng, Y., Bjorklund, A.K., Baque-Vidal, L., Sokka, J., Trokovic, R., et al. (2025). A comprehensive human embryo reference tool using single-cell RNA-sequencing data. Nat Methods 22, 193–206. 10.1038/s41592-024-02493-2.

21. Yagi, R., Kohn, M.J., Karavanova, I., Kaneko, K.J., Vullhorst, D., DePamphilis, M.L., and Buonanno, A. (2007). Transcription factor TEAD4 specifies the trophectoderm lineage at the beginning of mammalian development. Development 134, 3827–3836. 10.1242/dev.010223.

22. Nishioka, N., Inoue, K., Adachi, K., Kiyonari, H., Ota, M., Ralston, A., Yabuta, N., Hirahara, S., Stephenson, R.O., Ogonuki, N., et al. (2009). The Hippo signaling pathway components Lats and Yap pattern Tead4 activity to distinguish mouse trophectoderm from inner cell mass. Dev. Cell 16, 398–410. 10.1016/j.devcel.2009.02.003.

23. Hirate, Y., Hirahara, S., Inoue, K., Suzuki, A., Alarcon, V.B., Akimoto, K., Hirai, T., Hara, T., Adachi, M., Chida, K., et al. (2013). Polarity-Dependent Distribution of Angiomotin Localizes Hippo Signaling in Preimplantation Embryos. Curr. Biol. 23, 1181–1194. 10.1016/j.cub.2013.05.014.

24. Cockburn, K., Biechele, S., Garner, J., and Rossant, J. (2013). The Hippo Pathway Member Nf2 Is Required for Inner Cell Mass Specification. Curr. Biol. 23, 1195–1201. 10.1016/j.cub.2013.05.044.

25. Posfai, E., Petropoulos, S., de Barros, F.R.O., Schell, J.P., Jurisica, I., Sandberg, R., Lanner, F., and Rossant, J. (2017). Position-and Hippo signaling-dependent plasticity during lineage segregation in the early mouse embryo. Elife 6. 10.7554/eLife.22906.

26. Lorthongpanich, C., Messerschmidt, D.M., Chan, S.W., Hong, W., Knowles, B.B., and Solter, D. (2013). Temporal reduction of LATS kinases in the early preimplantation embryo prevents ICM lineage differentiation. Genes Dev. 27, 1441–1446. 10.1101/gad.219618.113.

27. Gerri, C., McCarthy, A., Mei Scott, G., Regin, M., Stamatiadis, P., Brumm, S., Simon, C.S., Lee, J., Montesinos, C., Hassitt, C., et al. (2023). A conserved role of the Hippo signalling pathway in initiation of the first lineage specification event across mammals. Development 150. 10.1242/dev.201112.

28. Gerri, C., McCarthy, A., Alanis-Lobato, G., Demtschenko, A., Bruneau, A., Loubersac, S., Fogarty, N.M.E., Hampshire, D., Elder, K., Snell, P., et al. (2020). Initiation of a conserved trophectoderm program in human, cow and mouse embryos. Nature 587, 443–447. 10.1038/s41586-020-2759-x.

29. Canizo, J.R., Zhao, C., and Petropoulos, S. (2025). The guinea pig serves as an alternative model to study human preimplantation development. Nat. Cell Biol. 27, 696–710. 10.1038/s41556-025-01642-9.

30. Kagawa, H., Javali, A., Khoei, H.H., Sommer, T.M., Sestini, G., Novatchkova, M., Scholte Op Reimer, Y., Castel, G., Bruneau, A., Maenhoudt, N., et al. (2021). Human blastoids model blastocyst development and implantation. Nature. 10.1038/s41586-021-04267-8.

31. Oura, S., Li, L., and Wu, J. (2025). An inducible model of human post-implantation development derived from primed and naive stem cells. Cell Stem Cell 32, 1509-1527.e9. 10.1016/j.stem.2025.08.005.

32. Pastor, W.A., Chen, D., Liu, W., Kim, R., Sahakyan, A., Lukianchikov, A., Plath, K., Jacobsen, S.E., and Clark, A.T. (2016). Naive Human Pluripotent Cells Feature a Methylation Landscape Devoid of Blastocyst or Germline Memory. Cell Stem Cell 18, 323–329. 10.1016/j.stem.2016.01.019.

33. Fischer, L.A., Meyer, B., Reyes, M., Zemke, J.E., Harrison, J.K., Park, K.M., Wang, T., Juppner, H., Dietmann, S., and Theunissen, T.W. (2025). Tracking and mitigating imprint erasure during induction of naive human pluripotency at single-cell resolution. Stem Cell Rep. 20, 102419. 10.1016/j.stemcr.2025.102419.

34. Kilens, S., Meistermann, D., Moreno, D., Chariau, C., Gaignerie, A., Reignier, A., Lelievre, Y., Casanova, M., Vallot, C., Nedellec, S., et al. (2018). Parallel derivation of isogenic human primed and naive induced pluripotent stem cells. Nat Commun 9, 360. 10.1038/s41467017-02107-w.

35. Azevedo Portilho, N., Saini, D., Hossain, I., Sirois, J., Moraes, C., and Pastor, W.A. (2021). The DNMT1 inhibitor GSK-3484862 mediates global demethylation in murine embryonic stem cells. Epigenetics Chromatin 14, 56. 10.1186/s13072-021-00429-0.

36. Zijlmans, D.W., Talon, I., Verhelst, S., Bendall, A., Van Nerum, K., Javali, A., Malcolm, A.A., van Knippenberg, S.S.F.A., Biggins, L., To, S.K., et al. (2022). Integrated multi-omics reveal polycomb repressive complex 2 restricts human trophoblast induction. Nat. Cell Biol. 24, 858–871. 10.1038/s41556-022-00932-w.

37. Kumar, B., Navarro, C., Winblad, N., Schell, J.P., Zhao, C., Weltner, J., Baque-Vidal, L., Salazar Mantero, A., Petropoulos, S., Lanner, F., et al. (2022). Polycomb repressive complex 2 shields naive human pluripotent cells from trophectoderm differentiation. Nat Cell Biol 24, 845–857. 10.1038/s41556-022-00916-w.

38. Theunissen, T.W., Friedli, M., He, Y., Planet, E., O’Neil, R.C., Markoulaki, S., Pontis, J., Wang, H., Iouranova, A., Imbeault, M., et al. (2016). Molecular Criteria for Defining the Naive Human Pluripotent State. Cell Stem Cell 19, 502–515. 10.1016/j.stem.2016.06.011.

39. Zhu, M., Meglicki, M., Lamba, A., Wang, P., Royer, C., Turner, K., Jauhar, M.A., Jones, C., Child, T., Coward, K., et al. (2024). Tead4 and Tfap2c generate bipotency and a bistable switch in totipotent embryos to promote robust lineage diversification. Nat. Struct. Mol. Biol. 31, 964–976. 10.1038/s41594-024-01311-9.

40. Piccolo, F.M., Kastan, N.R., Haremaki, T., Tian, Q., Laundos, T.L., De Santis, R., Beaudoin, A.J., Carroll, T.S., Luo, J.-D., Gnedeva, K., et al. (2022). Role of YAP in early ectodermal specification and a Huntington’s Disease model of human neurulation. eLife 11, e73075. 10.7554/eLife.73075.

41. Lee, C.Q., Gardner, L., Turco, M., Zhao, N., Murray, M.J., Coleman, N., Rossant, J., Hemberger, M., and Moffett, A. (2016). What Is Trophoblast? A Combination of Criteria Define Human First-Trimester Trophoblast. Stem Cell Rep. 6, 257–272. 10.1016/j.stemcr.2016.01.006.

42. Kobayashi, N., Okae, H., Hiura, H., Kubota, N., Kobayashi, E.H., Shibata, S., Oike, A., Hori, T., Kikutake, C., Hamada, H., et al. (2022). The microRNA cluster C19MC confers differentiation potential into trophoblast lineages upon human pluripotent stem cells. Nat. Commun. 13, 3071. 10.1038/s41467-022-30775-w.

43. Faure, L., Soldatov, R., Kharchenko, P.V., and Adameyko, I. (2023). scFates: a scalable python package for advanced pseudotime and bifurcation analysis from single-cell data. Bioinformatics 39. 10.1093/bioinformatics/btac746.

44. Liu, D.D., Chen, Y.D., Ren, Y.X., Yuan, P., Wang, N., Liu, Q., Yang, C., Yan, Z.Q., Yang, M., Wang, J., et al. (2022). Primary specification of blastocyst trophectoderm by scRNA-seq: New insights into embryo implantation. Sci Adv 8. https://doi.org/ARTN%2520eabj3725%252010.1126/sciadv.abj3725.

45. Meistermann, D., Bruneau, A., Loubersac, S., Reignier, A., Firmin, J., Francois-Campion, V., Kilens, S., Lelievre, Y., Lammers, J., Feyeux, M., et al. (2021). Integrated pseudotime analysis of human pre-implantation embryo single-cell transcriptomes reveals the dynamics of lineage specification. Cell Stem Cell 28, 1625–1640 e6. 10.1016/j.stem.2021.04.027.

46. Biondic, S., Cheng, Z., Yin, R., Theunissen, T., and Petropoulos, S. (2025). Dissecting Small Noncoding RNA Landscapes in Mouse Preimplantation Embryos and Human Blastoids for Modeling Early Human Embryogenesis. Preprint at bioRxiv, 10.1101/2025.10.07.680944 10.1101/2025.10.07.680944.

47. Vento-Tormo, R., Efremova, M., Botting, R.A., Turco, M.Y., Vento-Tormo, M., Meyer, K.B., Park, J.-E., Stephenson, E., Polański, K., Goncalves, A., et al. (2018). Single-cell reconstruction of the early maternal–fetal interface in humans. Nature 563, 347–353. 10.1038/s41586018-0698-6.

48. Petropoulos, S., Edsgard, D., Reinius, B., Deng, Q., Panula, S.P., Codeluppi, S., Reyes, A.P., Linnarsson, S., Sandberg, R., and Lanner, F. (2016). Single-Cell RNA-Seq Reveals Lineage and X Chromosome Dynamics in Human Preimplantation Embryos. Cell 167, 285. 10.1016/j.cell.2016.08.009.

49. Tyser, R.C.V., Mahammadov, E., Nakanoh, S., Vallier, L., Scialdone, A., and Srinivas, S. (2021). Single-cell transcriptomic characterization of a gastrulating human embryo. Nature 600, 285– 289. 10.1038/s41586-021-04158-y.

50. Yan, L., Yang, M., Guo, H., Yang, L., Wu, J., Li, R., Liu, P., Lian, Y., Zheng, X., Yan, J., et al. (2013). Single-cell RNA-Seq profiling of human preimplantation embryos and embryonic stem cells. Nat Struct Mol Biol 20, 1131–1139. 10.1038/nsmb.2660.

51. Xiang, L., Yin, Y., Zheng, Y., Ma, Y., Li, Y., Zhao, Z., Guo, J., Ai, Z., Niu, Y., Duan, K., et al. (2020). A developmental landscape of 3D-cultured human pre-gastrulation embryos. Nature 577, 537–542. 10.1038/s41586-019-1875-y.

52. Simunovic, M., Siggia, E.D., and Brivanlou, A.H. (2022). In vitro attachment and symmetry breaking of a human embryo model assembled from primed embryonic stem cells. Cell Stem Cell 29, 962-972.e4. 10.1016/j.stem.2022.05.001.

53. Arutyunyan, A., Roberts, K., Troule, K., Wong, F.C.K., Sheridan, M.A., Kats, I., Garcia-Alonso, L., Velten, B., Hoo, R., Ruiz-Morales, E.R., et al. (2023). Spatial multiomics map of trophoblast development in early pregnancy. Nature 616, 143–151. 10.1038/s41586-023-05869-0.

54. Yang, L.(杨利恒), Liang, P., Yang, H., and Coyne, C.B. (2023). Trophoblast organoids with physiological polarity model placental structure and function. J. Cell Sci. 137, jcs261528. 10.1242/jcs.261528.

55. Lee, C.Q.E., Turco, M.Y., Gardner, L., Simons, B.D., Hemberger, M., and Moffett, A. (2018). Integrin alpha2 marks a niche of trophoblast progenitor cells in first trimester human placenta. Development 145. 10.1242/dev.162305.

56. Shannon, M.J., McNeill, G.L., Koksal, B., Baltayeva, J., Wächter, J., Castellana, B., Peñaherrera, M.S., Robinson, W.P., Leung, P.C.K., and Beristain, A.G. (2024). Single-cell assessment of primary and stem cell-derived human trophoblast organoids as placenta-modeling platforms. Dev. Cell 59. 10.1016/j.devcel.2024.01.023.

57. Haider, S., Meinhardt, G., Saleh, L., Fiala, C., Pollheimer, J., and Knöfler, M. (2016). Notch1 controls development of the extravillous trophoblast lineage in the human placenta. Proc. Natl. Acad. Sci. U. S. A. 113, E7710–E7719. 10.1073/pnas.1612335113.

58. Skory, R.M. (2024). Revisiting trophectoderm-inner cell mass lineage segregation in the mammalian preimplantation embryo. Hum. Reprod. 39, 1889–1898. 10.1093/humrep/deae142.

59. Knofler, M., Haider, S., Saleh, L., Pollheimer, J., Gamage, T., and James, J. (2019). Human placenta and trophoblast development: key molecular mechanisms and model systems. Cell Mol Life Sci 76, 3479–3496. 10.1007/s00018-019-03104-6.

60. Dietrich, B., Haider, S., Meinhardt, G., Pollheimer, J., and Knofler, M. (2022). WNT and NOTCH signaling in human trophoblast development and differentiation. Cell Mol Life Sci 79, 292. 10.1007/s00018-022-04285-3.

61. Nehme, E., Panda, A., Migeotte, I., and Pasque, V. (2025). Extra-embryonic mesoderm during development and in in vitro models. Development 152, DEV204624. 10.1242/dev.204624.

62. Pham, T.X.A., Panda, A., Kagawa, H., To, S.K., Ertekin, C., Georgolopoulos, G., van Knippenberg, S.S.F.A., Allsop, R.N., Bruneau, A., Chui, J.S.-H., et al. (2022). Modeling human extraembryonic mesoderm cells using naive pluripotent stem cells. Cell Stem Cell 29, 1346-1365.e10. 10.1016/j.stem.2022.08.001.

63. Chen, C., Ridzon, D.A., Broomer, A.J., Zhou, Z., Lee, D.H., Nguyen, J.T., Barbisin, M., Xu, N.L., Mahuvakar, V.R., Andersen, M.R., et al. (2005). Real-time quantification of microRNAs by stemloop RT-PCR. Nucleic Acids Res. 33, e179. 10.1093/nar/gni178.

64. Peltier, H.J., and Latham, G.J. (2008). Normalization of microRNA expression levels in quantitative RT-PCR assays: Identification of suitable reference RNA targets in normal and cancerous human solid tissues. RNA 14, 844–852. 10.1261/rna.939908.

65. Kumaki, Y., Oda, M., and Okano, M. (2008). QUMA: quantification tool for methylation analysis. Nucleic Acids Res. 36, W170–W175. 10.1093/nar/gkn294.

66. Khan, S.A., Park, K.M., Fischer, L.A., Dong, C., Lungjangwa, T., Jimenez, M., Casalena, D., Chew, B., Dietmann, S., Auld, D.S., et al. (2021). Probing the signaling requirements for naive human pluripotency by high-throughput chemical screening. Cell Rep 35. https://doi.org/ARTN%2520109233%252010.1016/j.celrep.2021.109233.

67. Bredenkamp, N., Yang, J., Clarke, J., Stirparo, G.G., von Meyenn, F., Dietmann, S., Baker, D., Drummond, R., Ren, Y., Li, D., et al. (2019). Wnt Inhibition Facilitates RNA-Mediated Reprogramming of Human Somatic Cells to Naive Pluripotency. Stem Cell Rep. 13, 1083–1098. 10.1016/j.stemcr.2019.10.009.

68. Mazid, M.A., Ward, C., Luo, Z., Liu, C., Li, Y., Lai, Y., Wu, L., Li, J., Jia, W., Jiang, Y., et al. (2022). Rolling back human pluripotent stem cells to an eight-cell embryo-like stage. Nature 605, 315– 324. 10.1038/s41586-022-04625-0.

69. Yu, G., Wang, L.-G., Han, Y., and He, Q.-Y. (2012). clusterProfiler: an R Package for Comparing Biological Themes Among Gene Clusters. OMICS J. Integr. Biol. 16, 284–287. 10.1089/omi.2011.0118.

70. Taubenschmid-Stowers, J., Rostovskaya, M., Santos, F., Ljung, S., Argelaguet, R., Krueger, F., Nichols, J., and Reik, W. (2022). 8C-like cells capture the human zygotic genome activation program in vitro. Cell Stem Cell 29, 449–459 e6. 10.1016/j.stem.2022.01.014.

71. Kavaliauskaite, G., and Madsen, J.G.S. (2023). Automatic quality control of single-cell and single-nucleus RNA-seq using valiDrops. NAR Genomics Bioinforma. 5, qad101. 10.1093/nargab/lqad101.

72. Young, M.D., and Behjati, S. (2020). SoupX removes ambient RNA contamination from droplet-based single-cell RNA sequencing data. GigaScience 9, giaa151. 10.1093/gigascience/giaa151.

73. Germain, P.-L., Lun, A., Meixide, C.G., Macnair, W., and Robinson, M.D. (2022). Doublet identification in single-cell sequencing data using scDblFinder. Preprint at F1000Research, 10.12688/f1000research.73600.2 10.12688/f1000research.73600.2.

