## Supplemental Figures for "Reconstructing the emergence of the human chorion via HIPPO-mediated trophoblast induction"

### Supplementary Figures

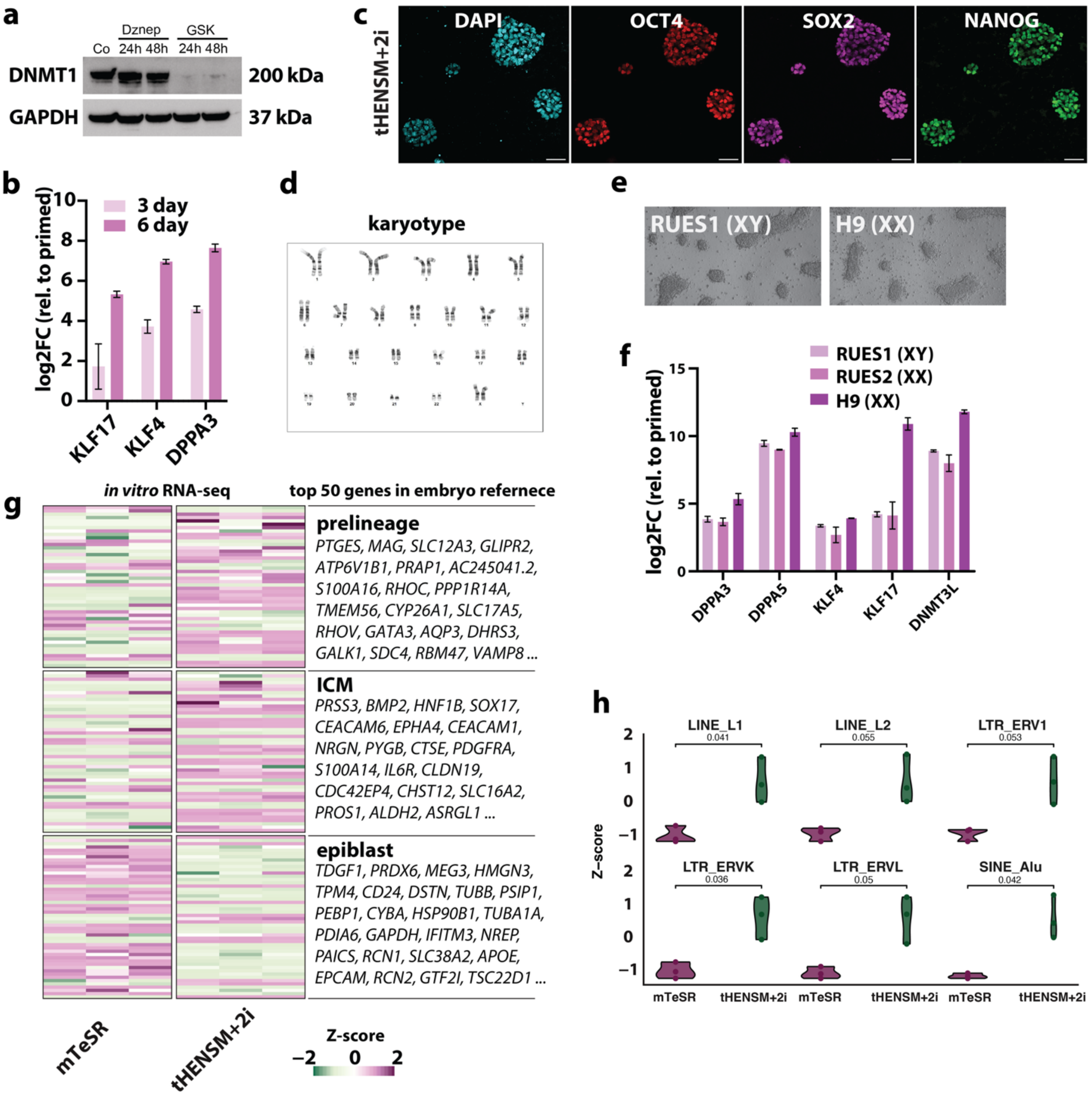


*Figure S1: Validation of transient naïve resetting across hESC lines.* *(a) Western blot showing depletion of DNMT1 protein after treatment with GSK-3484862. (b) qPCR analysis of naïve hESC genes after three and six days in HENSM + 2i relative to primed hESCs (n = 3; bars show mean ± s.d.). (c) IF for pluripotency markers in tHENSM+2i (six-day treatment). Scale bars, 50 µm. (d) Representative karyotype of RUES2 hESCs. (e-f) Bright-field images of additional hESC lines cultured in tHENSM+2i and qPCR analysis showing consistent induction of naïve genes across three lines (n = 3; mean ± s.d.). (g) Heatmap showing the expression of the 50 most differentially expressed genes by the prelineage, ICM, and late-epiblast clusters from the integrated human embryo reference (ref. 20) applied to our bulk RNA-seq data from mTeSR and tHENSM+2i cultures. (h) Violin plots showing Z-score-normalized expression of representative transposable elements in primed versus tHENSM+2i cultures.*

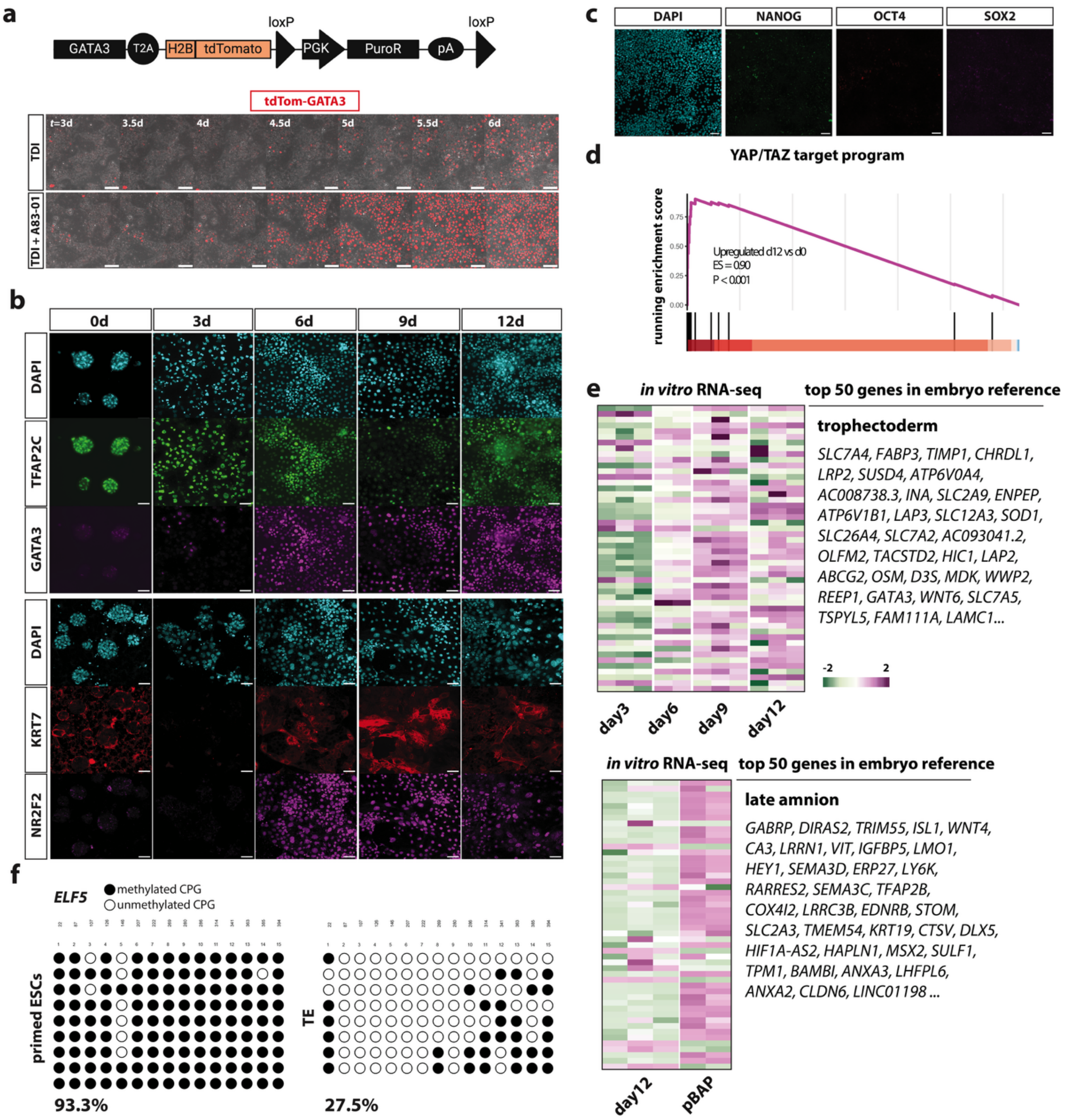


***Figure S2**: **HIPPO modulation induces a YAP/TEAD-dependent TE program.*** *(a) Schematic of the GATA3-tdTom reporter construct and live-cell time-lapse montages from the first six days under TDI alone or TDI + A83-01 conditions, showing progressive activation of the reporter in either condition. (b) IF time course showing progressive protein expression of canonical TE genes. Scale bars, 50 µm. (c) IF on day 12 shows absence of pluripotency markers. Scale bars, 50 µm. (d) Gene set enrichment analysis (GSEA) of a YAP/TAZ transcriptional target program on bulk RNA-seq (day 12 vs day 0), confirming transcriptional activation of canonical YAP targets. (e) Bisulfite sequencing of the ELF5 promoter (−136 to +238 bp relative to TSS). Each row represents a sequenced clone; filled and open circles denote methylated and unmethylated CpG sites, respectively. (f) Expression of the 50 most differentially expressed genes in the TE and amnion clusters calculated from the integrated embryonic reference (ref-20) applied to our in vitro RNA-seq time course. Top, expression dynamics of TE genes from day 0 to 12; bottom, comparison of late amnion genes between our day-12 cells and BAP-treated primed hESCs (data reproduced from ref-19). Values are Z-score scaled.*

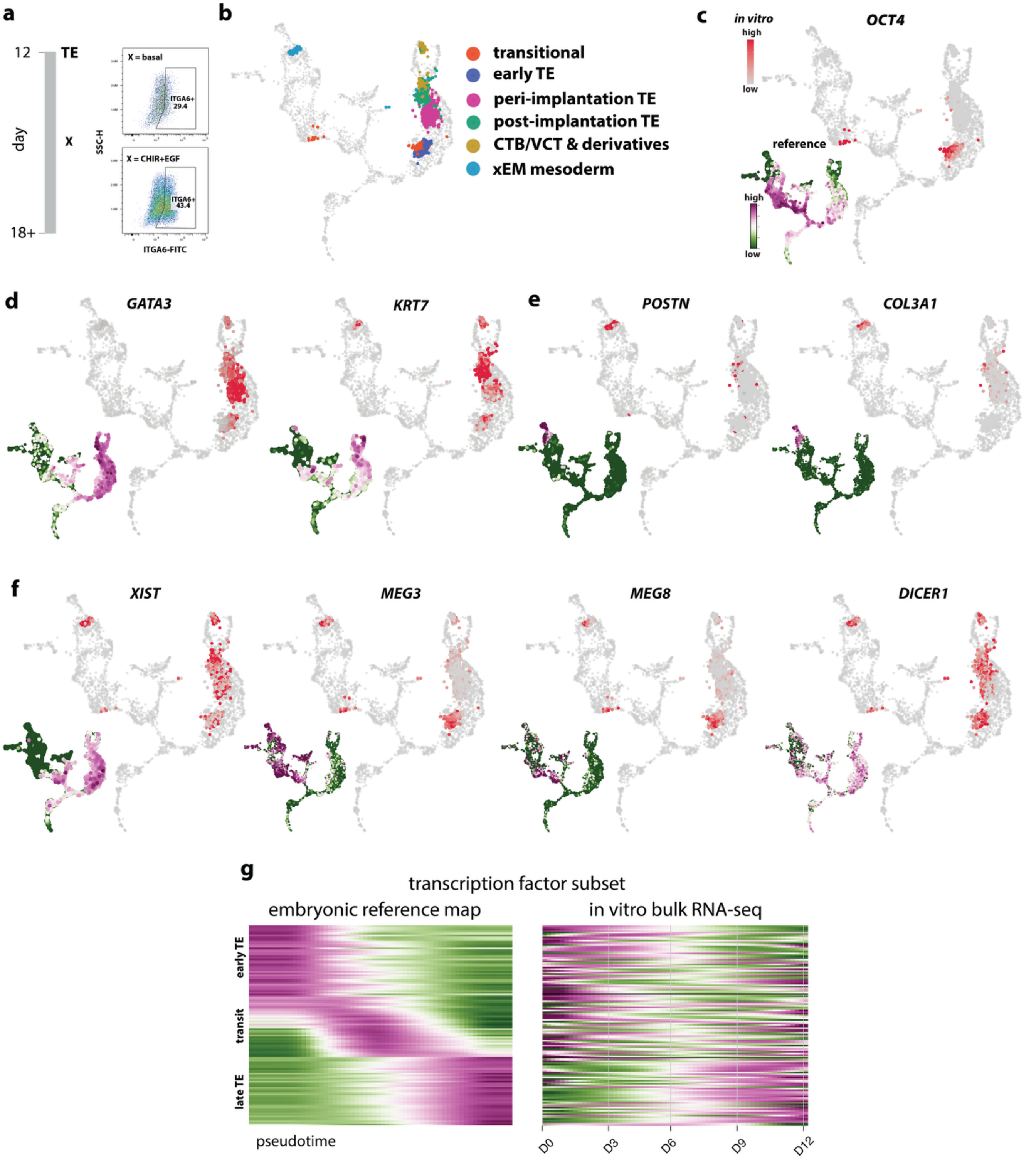


*__Figure S3: Further characterization of temporal dynamics downstream of HIPPO.__ (a) Flow cytometry for ITGA6 on day 18 under basal versus CHIR/EGF conditions. (b) Clustering of the combined projected day-6/12/18 in vitro single-cell datasets within the stabilized UMAP embedding of the integrated embryonic reference. (c-f) Gene expression overlays: large panels show expression in the projected in vitro datasets (reference cells shown in gray), while small insets display the corresponding expression domains in the embryonic reference. (g) Transcription-factor subset ordered along the embryonic TB pseudotime trajectory in the embryonic reference (left) and applied to the in vitro bulk RNA-seq time course (right). Showing a 10% subset of early, transit, and late TE modules for clarity.*

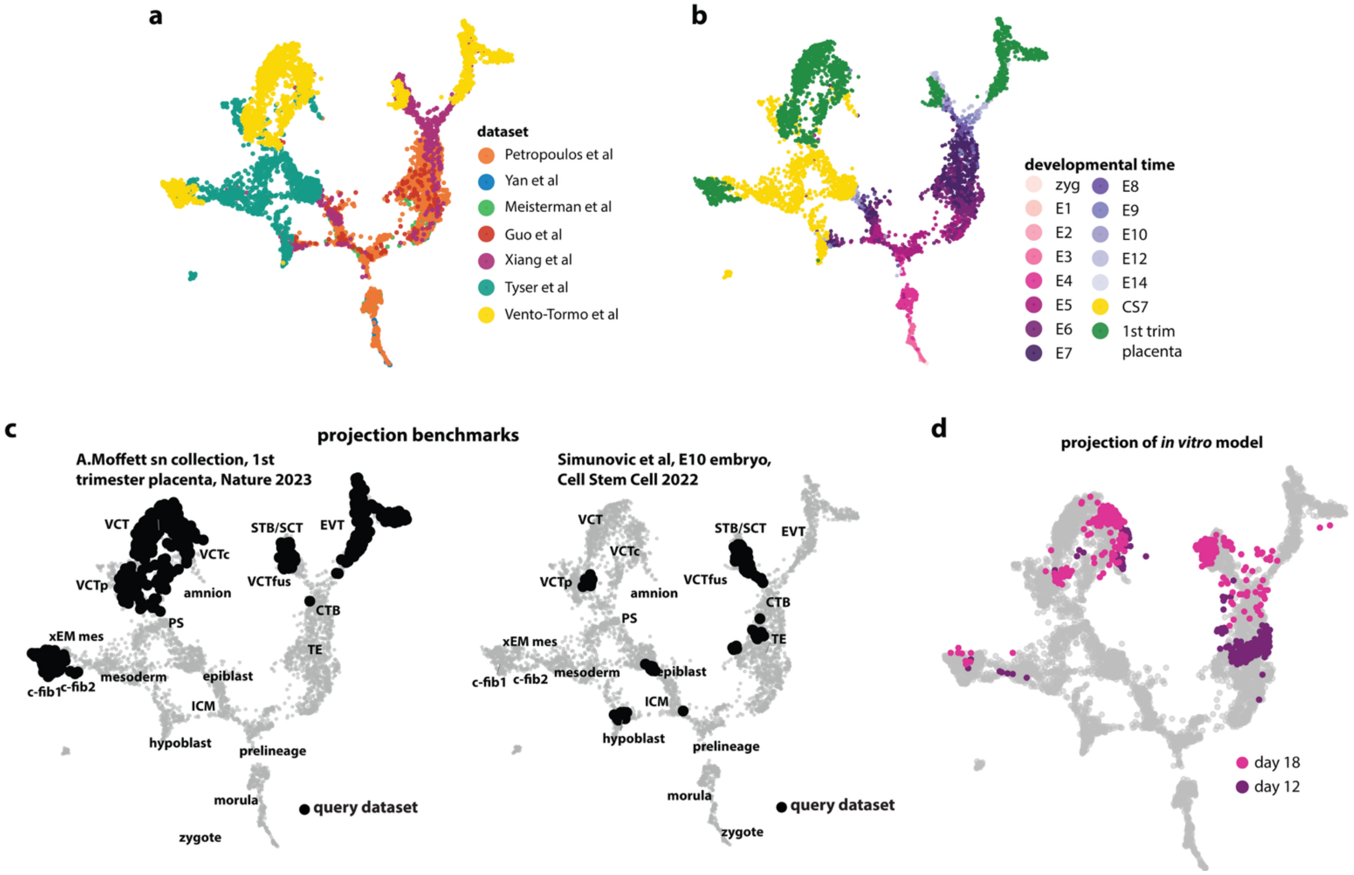


*Figure S4: Benchmarking the extended embryo-placenta reference and further projection data.* (a-b) UMAP representation of the extended reference map showing dataset composition (a) and developmental staging from zygote through first-trimester placenta (b). (c) Projection benchmarks of independent datasets not used for integration, including single-nuclear data from first-trimester placenta (the Ashley Moffett collection from ref-53) and E10 human embryo data (from ref-52), demonstrating correct stage and lineage alignment. (d) Projection of day-12 and day-18 in vitro transcriptomes onto the extended map, showing temporal ordering and representation across chorionic lineages.

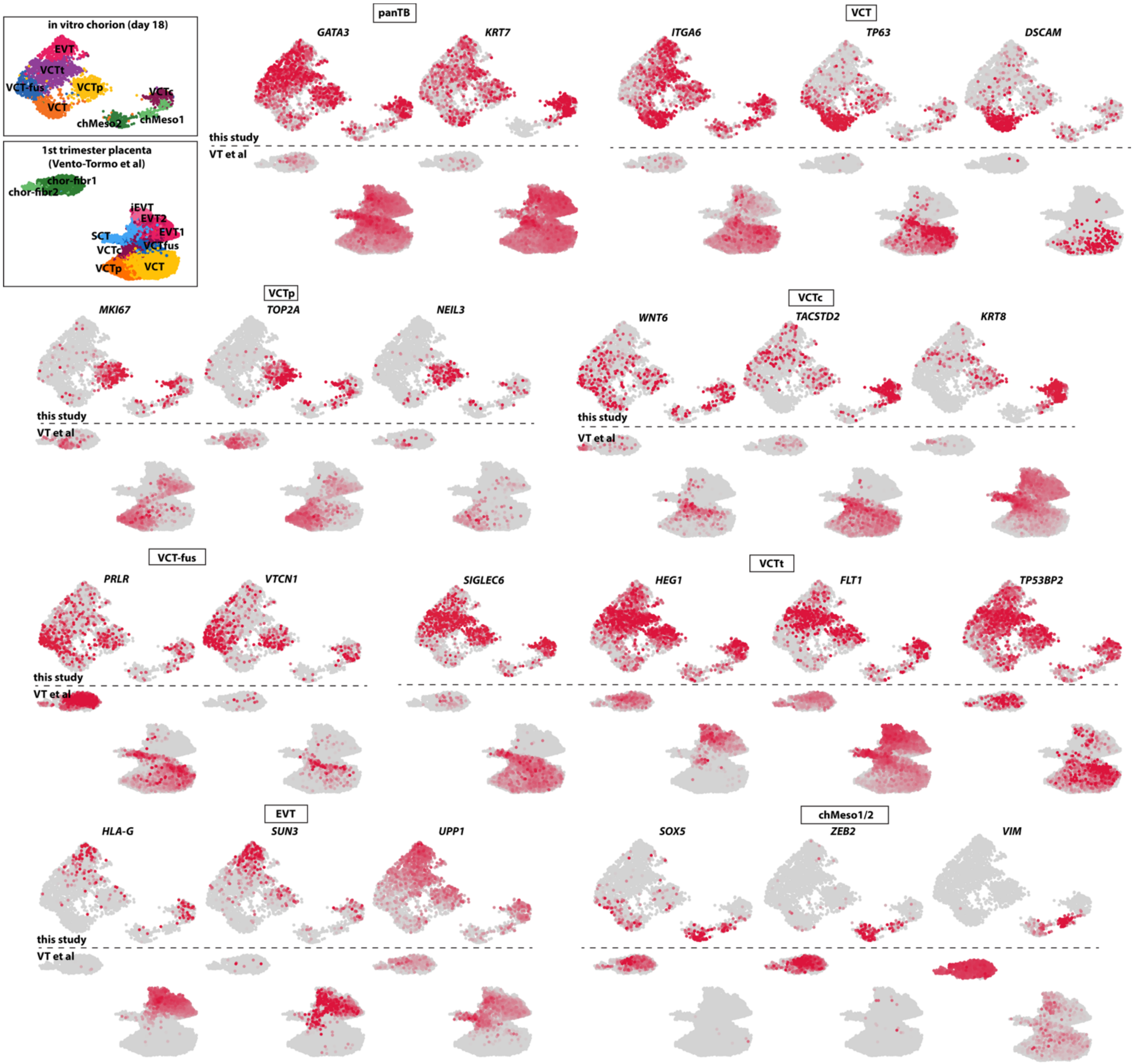


***Figure S5: Comparison of in-vitro day-18 chorion cultures with first-trimester placenta.*** *Inset (upper left) shows UMAP cluster assignments for the in-vitro day-18 chorion (top) and for the first-trimester placenta reference reproduced from data deposited by Vento-Tormo and colleagues, from ref-47 (bottom).*

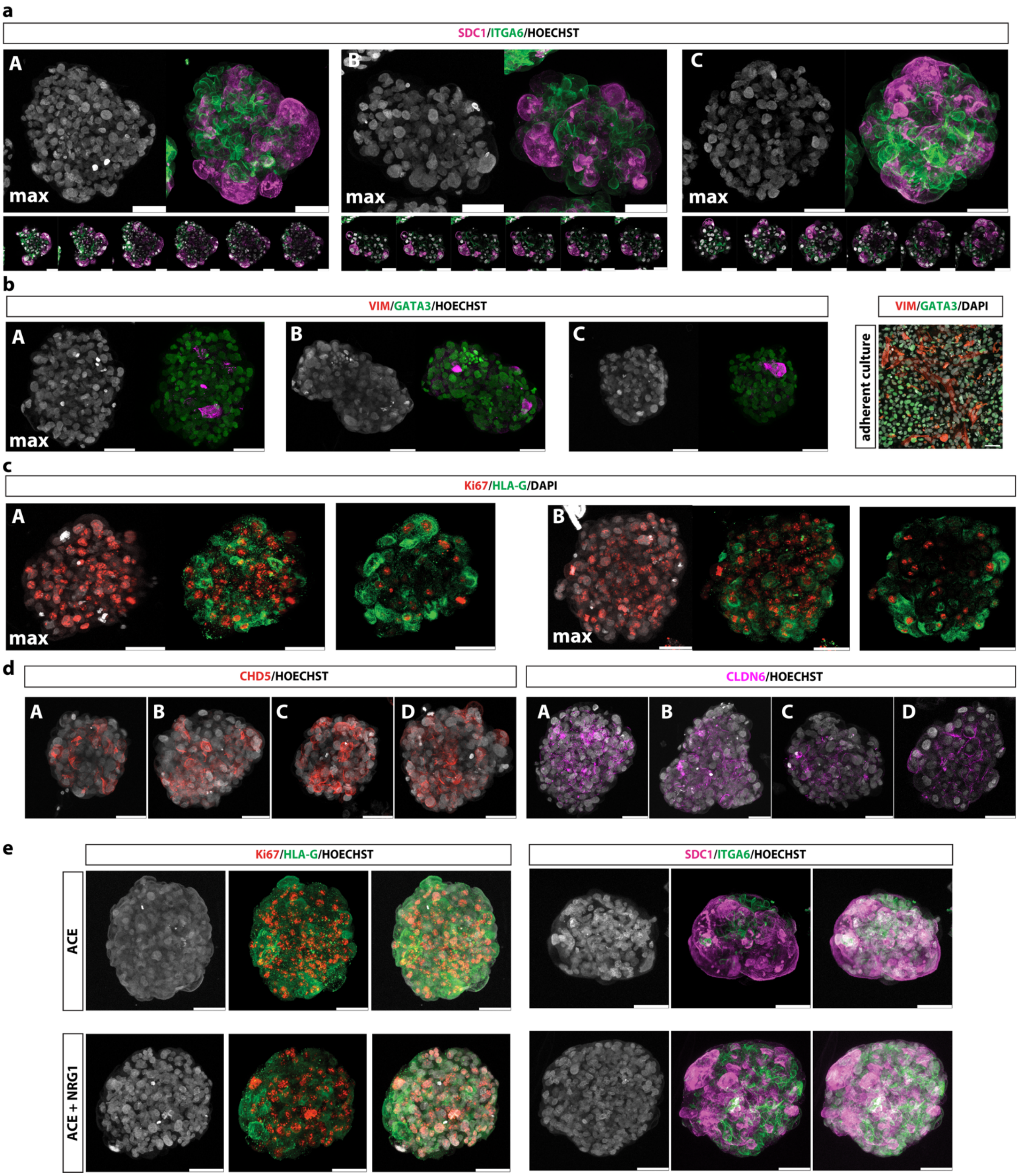


***Figure S6: Further examples of chorion organoids.*** *(a-d) IF stains of optically cleared chorion organoids, showing appropriate polarity (a), mesodermal-TB compartments (b) and EVT emergence (c, d). Each stain shows multiple independent examples (marked with capital letters). Max = maximum projection. Adherent culture in b is from day-18 experiment. (e) IF stains of chorion organoids under alternative culture conditions. ACE = A83-01, CHIR99021, EGF1; NRG1 = neuregulin 1. Scale bars, 50 µm.*
